# Nitrotoxin metabolism in bacteria may have emerged from a diverse oxidoreductase reservoir

**DOI:** 10.64898/2026.09.13.751228

**Authors:** Partha Barman, Shilpa Sinha, Ranadhir Chakraborty

**Affiliations:** OMICS Laboratory, Department of Biotechnology, University of North Bengal, Siliguri-734013, West Bengal, India

**Keywords:** 3-NPA, *Serratia*, Nitrotoxin, YrpB, Metagenomics, Transcriptome, MD simulation

## Abstract

Bacterial transformation of 3 nitropropionic acid (3NPA) is typically viewed through canonical nitronate monooxygenases (NMOs), yet sequence annotation poorly resolves function across this flavoprotein family. Here we show that the environmental YrpB/NMO associated oxidoreductase space is dominated by YrpB related proteins: across 13 metagenomes, 83.1% of 390 conserved domain supported candidates were YrpB-associated. Cultivation along an *Eisenia fetida* feed gut cast continuum recovered 31 phylogenetically diverse 3NPA responsive bacteria, revealing that this phenotype is distributed across multiple lineages. Using the previously genome-sequenced *Serratia* sp. EWG9 as a tractable exemplar, we demonstrate sustained 3NPA supported growth, 44% parent compound depletion and a broad early transcriptional reorganization. Its focal oxidoreductase OXR01 occupies overlapping YrpB/NMO sequence space and shows stable predicted 3NPA accommodation without strong transcriptional induction. These findings reposition canonical NMOs as one component of a broader, ecologically distributed oxidoreductase reservoir for bacterial nitrotoxin responsiveness.

## INTRODUCTION

Toxins from the food have a basic limitation for niche diversification in animals. It is especially true in relation to the toxins in the plants and microbes, such as 3-nitropropionic acid (3-NPA), which are abundant in plant forage and detrital material and which can irreversibly suppress essential metabolic functions. Still, animals manage to survive on such toxin-enriched diets in different environments. This paradox reveals a crucial importance of the microbiota as a biochemical bridge connecting the environment and host physiology. Diet-associated and gut microbiota may convert toxic substances into metabolizable forms or at least make them less toxic, thereby protecting host metabolism and gaining access to novel resources. However, the enzymatic nature of the process and whether it involves conserved or diverse biochemical pathways are unknown.

This interaction between microbes and the degradation of 3-NPA is a clear example of such an interface. Microorganisms capable of detoxifying and metabolizing 3-NPA have been found in herbivorous animals, some of which use 3-NPA as a special source of carbon, nitrogen, and energy^1,2^.

However, this ability is present within complex microbial communities. When plant material is ingested, the microbial communities undergo significant changes as they pass through the gut and are then transferred into the detrital food web, resulting in the development of interconnected microbiomes that are both associated with the host and found in the environment. As a result, cultivation of microorganisms can only reveal a small portion of the genetic material involved in nitrotoxin metabolism^3,4^. It therefore follows that these biological processes should be studied using both cultivation-based and cultivation-free methods.

The usual biochemical pathway for the metabolism of 3-NPA in bacteria is based on a group of flavin-dependent nitronate monooxygenases (NMOs) in the COG2070 family. The structures of these proteins are well understood and their kinetic properties have been determined; it is believed that they carry out the oxidation of propionate 3-nitronate and thus link the breakdown of the nitrotoxin to central metabolic pathways^5–7^. Yet new data raises questions about the completeness of this pathway. Functional annotations of NMOs are not always correct: some proteins identified as NMOs do not show monooxygenase activity, and some organisms that metabolize 3-NPA may use oxidoreductases which do not belong to the category of conventional NMOs. Hence, the question to be asked is whether the metabolism of nitrotoxins is controlled by the conserved enzymatic machinery or instead is the result of a wider range of oxidoreductases?

In addressing this question, we combine cultivation-based assessment of 3-NPA-responsive bacteria with a genome/metagenome-centric exploration through interconnected ecosystems. Comparison of several metagenomes associated with the gut, manure, and soil environment^8–10^ demonstrates that the proposed oxidoreductases are characterized not by the common NMO structural elements but by a widespread group of YrpB-associated oxidoreductases. Phylogenetic analysis shows that the NMO-like proteins belong to the broader sequence landscape suggesting that classical NMOs form a small and specialized clade inside the diverse oxidoreductase framework.

In accordance with this paradigm, phylogenetically distinct bacteria which were able to grow in 3-NPA-dependent way were identified along the *Eisenia fetida* food–digestive tract–casts axis; however, oxidoreductase enzymes of these bacteria were located in different regions of the sequence space, related to the YrpB and NMO enzymes. Here we describe a strain of *Serratia* sp., which was grown in 3-NPA-dependent way, depletion of the initial compound, and the existence of YrpB-related enzyme of partially similar structure to NMO. However, analysis of transcriptome data demonstrates that such approaches are not sufficient for functional assignment of enzymes.

Collectively, these observations place bacterial nitrotoxin responsiveness within a broader YrpB/NMO-associated oxidoreductase sequence space than is captured by canonical NMO annotation alone. This framework identifies an ecologically widespread reservoir of candidate flavin oxidoreductases whose contributions to nitrotoxin transformation remain to be resolved experimentally.

## ONLINE METHODS

### Study design and analytical framework

This study used a hierarchical analytical framework to examine the ecological distribution, sequence diversity and functional context of YrpB- and nitronate monooxygenase (NMO)-associated oxidoreductases. We first surveyed independently generated whole-metagenome datasets representing contrasting ecological environments to determine the distribution and relative representation of YrpB-associated and NMO-like protein architectures. Candidate proteins were subsequently examined using conserved-domain and phylogenetic analyses to define the broader sequence space within which the focal *Serratia* sp. EWG9 oxidoreductase occurs.

The environmental analysis was then connected to an experimentally accessible *Eisenia fetida* feed– gut–cast system. Cultivable bacteria capable of growth under 3-nitropropionic acid (3-NPA)-associated conditions were recovered from this continuum and characterized by 16S rRNA gene sequencing. Genome-resolved representatives were compared for their YrpB/NMO-associated protein repertoires and growth phenotypes, leading to prioritization of *Serratia* sp. EWG9 for detailed physiological, chemical, transcriptomic and structural investigation. The final analytical layer comprised molecular docking, protein–ligand interaction analysis, 300-ns molecular-dynamics simulations and binding-pocket comparison of the focal EWG9 protein and reference/comparator proteins.

Environmental metagenomes represented independently sourced datasets rather than biological replicates. Ecological grouping of these datasets was therefore used for comparative visualization and descriptive interpretation and not for replicate-based inferential testing.

### Environmental metagenome selection, YrpB/NMO candidate retrieval and quantitative analysis

Thirteen independently generated whole-metagenome datasets were selected to survey the distribution of YrpB/NMO-associated protein architectures across four broad ecological contexts: earthworm- or compost-associated environments, herbivore- or host-associated environments, agricultural or soil environments, and a waste-associated environment. The datasets comprised an earthworm gut metagenome (EGM), NBU vermicompost metagenome (NBUVC), soil amended with raw cow manure (SMWCM), bovine faecal metagenome (BFM), bovine rumen metagenome (BRM), two buffalo rumen metagenomes (BR1M and BR2M), chicken gastrointestinal tract metagenome (CGIT), farmland soil from the USA (FSUS), two rice-field soil metagenomes from Odisha (RF1O and RF2O), cotton rhizosphere soil from Rajasthan (CRSR), and top landfill soil from Nagpur (TLSN). Dataset metadata and accession numbers are provided in Supplementary Table 2. The datasets were selected to provide representation across contrasting ecological settings and were not treated as biological replicates of the corresponding ecological categories.

Whole-metagenome FASTA files were retrieved from the NCBI Genome database, and each dataset was processed independently using the same analytical workflow. Functional annotation was performed on the Galaxy Europe server^11^ using eggNOG-mapper v2.1.13 with the eggNOG v5.0.2^12^ database in metagenomic mode and DIAMOND in sensitive mode. Predicted protein sequences from each metagenome were subsequently screened for conserved-domain correspondence to three domain models representing the broader YrpB/NMO-associated protein space: YrpB/COG2070, NPD_like/cd04730 and NMO/PF03060.

Candidate discovery was performed independently of protein annotation terminology. Proteins showing correspondence to at least one of the COG2070/YrpB, cd04730/NPD_like or PF03060/NMO domain models were retained as the initial candidate pool. Corresponding protein sequences were retrieved and examined individually using NCBI Conserved Domain Search^13–15^ to determine their complete conserved-domain architecture and the relative strength of correspondence to the three domain models.

To provide an additional sequence-based criterion independent of conserved-domain annotation, candidate proteins were compared with selected reference proteins representing experimentally characterized nitronate monooxygenases and functionally characterized or representative YrpB-family oxidoreductases. Reference-sequence similarity, conserved-domain architecture and phylogenetic placement were evaluated together during candidate classification.

Candidates were operationally classified into two sequence-based groups. Proteins showing predominant correspondence to COG2070/YrpB and lacking stronger NMO-associated domain or phylogenetic support were classified as YrpB-associated. Proteins showing strong correspondence to PF03060/NMO and/or cd04730/NPD_like together with placement within or adjacent to experimentally characterized NMO-associated sequence groups were classified as NMO-associated. Proteins showing substantial overlap between YrpB-, NPD_like- and NMO-associated domain models, without sufficient evidence for reliable separation, were sorted according to their top domain architecture. Classification therefore incorporated conserved-domain architecture, reference-protein similarity and phylogenetic context rather than annotation terminology or a single domain hit alone. These operational categories describe sequence and domain relationships and were not interpreted as evidence of experimentally demonstrated enzymatic activity or substrate specificity.

For each metagenome, raw numbers of YrpB-associated and NMO-associated candidates were recorded. Because the total number of annotated proteins differed among datasets, candidate representation was additionally expressed relative to the annotated protein complement of the corresponding metagenome:

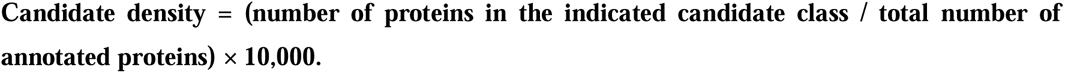

Candidate density therefore represents the number of conserved-domain-verified candidates per 10,000 annotated proteins and was used to account partially for differences in annotated protein-set size among datasets. This metric was not interpreted as a sequencing-depth-normalized estimate of gene, organismal or community abundance.

The proportional representation of each candidate class within the total retrieved YrpB/NMO-associated candidate set was calculated as:

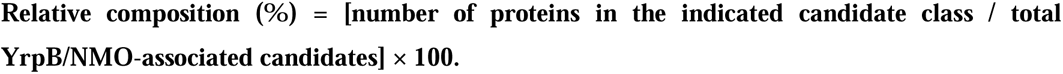

Raw candidate counts, candidate density and relative composition were used as complementary descriptive measures. Raw counts represented the number of candidate proteins recovered from each dataset, candidate density described their representation relative to the annotated protein complement, and relative composition described the proportional contribution of the operational candidate classes within the retrieved YrpB/NMO-associated sequence pool.

Because the metagenomes originated from independent studies and differed in sampling, sequencing and assembly backgrounds, ecological categories were used only for descriptive comparison and visualization. Individual datasets were not treated as biological replicates, and replicate-based inferential statistical tests were not applied across ecological categories.

For phylogenetic analysis, all conserved-domain-verified candidates recovered from the 13 environmental metagenomes were combined with selected reference proteins representing the broader YrpB/NMO-associated sequence space. Experimentally characterized proteins were included as functional anchors where available. These included PA4202 from *Pseudomonas aeruginosa*, an experimentally characterized Class I nitronate monooxygenase^5^, and PA1024, an experimentally characterized NADH:quinone reductase representing a non-NMO YrpB-family function^6^. Additional functionally unresolved YrpB/NMO-associated homologues were included to provide broader sequence context.

Amino-acid sequences were aligned using MUSCLE^16^, and maximum-likelihood phylogenetic reconstruction was performed in MEGA^17^ using the Jones–Taylor–Thornton substitution model.

Branch support was evaluated using 1,000 bootstrap replicates^18^. Phylogenetic placement was interpreted together with conserved-domain architecture and reference-protein similarity to assess the sequence relationships of environmental candidates. Proximity to an experimentally characterized reference protein was not considered independently sufficient for assigning enzymatic function.

### Establishment of the *Eisenia fetida* feed–gut–cast system, isolation and taxonomic characterization of cultivable 3-NPA-responsive bacteria

The *Eisenia fetida* feed–gut–cast system was established as described previously^19^. Raw cow dung (RCD) was processed to obtain an odour-reduced substrate for earthworm maintenance, and healthy adult E. fetida were maintained in processed RCD (PrCD) at 22 °C and 70–80% relative humidity. For preparation of the formulated feed, fresh *Quisqualis indica* bark and leaves were mixed at a 3:2 ratio, and approximately 250 g of plant material was incorporated between layers of approximately 1,250 g RCD and composted for 3–4 weeks under shaded conditions with periodic moistening using sterile deionized water. For the experimental setup, 75 g of composted formulated feed was placed in sterile 14-cm Petri plates, and each plate received six clitellated *E. fetida* weighing 0.30 ± 0.05 g each. Plates were maintained in the dark at 27 °C and periodically moistened with sterile deionized water. A corresponding PrCD-fed system was maintained as the normal-feed control.

Cultivable bacteria were recovered from formulated feed, whole gut of formulated-feed-fed earthworms and freshly produced casts, together with samples from the corresponding normal-feed control continuum. Approximately 0.1 g of feed or cast material was suspended in sterile phosphate-buffered saline (PBS). For gut-associated recovery, earthworms were washed thoroughly to remove externally adhering material, aseptically dissected, and the excised whole gut with its contents was macerated in sterile PBS. The resulting suspensions were serially diluted and spread onto agar fortified minimal salt medium (MSM) containing glucose and 3-nitropropionic acid (3-NPA; 0.06 g l^−1^), with 3-NPA replacing inorganic nitrate as the sole nitrogen source. Plates were incubated at 30□°C for 2–3 d, or until discrete colonies appeared. Morphologically distinct colonies were purified by repeated streaking on the same selective medium and reassessed for reproducible growth in glucose– 3-NPA MSM containing 0.06 g l^−1^ 3-NPA. Thirty-one isolates showing consistent growth under these conditions were retained for further analysis. Isolates recovered from formulated feed, whole gut and casts were designated MLF, EWG and EMC, respectively, whereas those from the normal-feed control continuum were designated PR. Because selection was based on reproducible growth under 3-NPA-containing conditions rather than direct measurement of substrate disappearance, these strains are referred to as 3-NPA-responsive isolates.

Genomic DNA from the 31 isolates was used to amplify the nearly full-length 16S rRNA gene using the universal bacterial primer pair 27F and 1492R. Amplicons were subjected to Sanger sequencing, and consensus sequences were compared against the NCBI nucleotide database using BLASTn for taxonomic identification. Sequences were aligned in MEGA12^17^ and used to construct a maximum-likelihood phylogenetic tree with 1,000 bootstrap replicates^18^. The same 16S rRNA gene sequences were also subjected to *in silico Hae*III digestion, and the predicted restriction fragments were visualized as virtual gel profiles. Because these restriction profiles were derived from the same 16S rRNA sequences used for phylogenetic reconstruction, they were treated as a complementary representation of sequence-level diversity rather than as an independent phylogenetic dataset. Sequence accession numbers are provided in Supplementary Table 3.

### Culture–amplicon metagenome genus-level taxonomic correspondence

To determine the extent to which selective cultivation recovered bacterial lineages represented in the corresponding culture-independent community, genus-level identities of the cultivable 3-NPA-responsive isolates along with earlier legacy isolates were compared with bacterial genera detected in the 16S rRNA gene amplicon datasets from the *Eisenia fetida* feed–gut–cast system. The cultivable collection comprised isolates recovered from the present *Quisqualis indica*-amended feed–gut–cast continuum together with legacy isolates obtained previously from the conventional PrCD-fed *E. fetida* feed–gut–cast system. The latter therefore provided additional representation of the unamended control continuum.

For the culture-independent comparison, previously generated amplicon datasets representing PrCD, formulated feed, earthworm gut and cast/vermicompost-associated samples were used. These datasets were generated by V3-region 16S rRNA gene sequencing, with taxonomic assignments obtained using the RDP Classifier at an 80% confidence threshold^20^. Genus-level assignments from the current and legacy cultivable collections were standardized and compared with the corresponding amplicon-derived genus inventory to identify shared and method-specific taxa. The comparison was used as a qualitative assessment of taxonomic correspondence between culture-dependent recovery and culture-independent detection, rather than as a quantitative comparison of relative abundance.

### Comparative growth phenotyping of cultivable 3-NPA-responsive bacteria

To compare the relative growth performance of the cultivable collection under 3-NPA-associated conditions, the 31 *E. fetida*-associated isolates were grown in glucose-supplemented MSM containing 3-NPA (0.06 g l^−1^) as the sole nitrogen source. Growth was followed over time, and generation times were calculated from several different timepoints of the exponential-growth phase of the corresponding growth profiles using g = (t_2_−t_1_) log_10_ 2/(log_10_N2−log_10_N1), with values calculated independently for each biological replicate. The mean generation-time comparison was used as a phenotypic screen to identify rapidly growing representatives within the broader cultivable collection.

On the basis of this screening, *Serratia* sp. EWG9 was identified as one of the fastest-growing 3-NPA-responsive isolates, whose whole-genome sequence had been reported previously and was therefore included in a focused comparative analysis together with selected genome-resolved legacy isolates previously recovered from the PrCD-fed *E. fetida* feed–gut–cast system^4^. In parallel, optimum temperature and optimum pH for growth for strain EWG9 were determined in glucose-supplemented MSM containing 3-NPA (0.06 g l^−1^) as the sole nitrogen source (n = 3). These included *Mycobacterium* sp. EPG1, *Streptomyces* sp. EAG2 and *Pradoshia eiseniae* EAG3, whose available genome annotations contained putative nitronate-monooxygenases^4^. *Pseudomonas aeruginosa* CD3, a soil-derived laboratory isolate^21^, and *Bacillus subtilis* MTCC121 were included as additional comparator strains representing *Pseudomonas*- and *Bacillus*-associated NMO/YrpB protein backgrounds, respectively.

For comparative growth analysis, fresh cultures derived from purified single colonies were grown in Luria broth, and 1% (v/v) of each culture was transferred into glucose–KNO_3_ MSM for preparation of the mother inoculum. Cultures were grown to mid-logarithmic phase, after which 1% (v/v) of the mother inoculum was used to inoculate glucose–3-NPA MSM containing 0.06 g l^−1^ 3-NPA. Cultures were incubated at 30□°C with shaking at 100 rpm, and growth was monitored at 12-h intervals for 72 h by viable-cell enumeration following serial dilution and plating on agar-solidified Luria broth medium. Viable counts were expressed as log_10_ CFU ml^−1^. Experiments were performed using three independent biological replicates.

### Genome-based retrieval and preliminary identification of YrpB/NMO-associated proteins

Whole-genome sequence data from the selected 3-NPA-responsive representatives were compiled for comparative analysis, principally *Serratia* sp. EWG9, *Streptomyces* sp. EAG2, *Mycobacterium* sp. EPG1 and *Pradoshia eiseniae* EAG3. Corresponding proteins recovered from the *Eisenia fetida* gut metagenome and genome-resolved MAGs were also included where required.

Predicted protein-coding sequences were screened using available genome annotations, RAST and orthology- and domain-based searches. Proteins annotated as nitronate monooxygenases, 2-nitropropane dioxygenases, YrpB-family oxidoreductases, nitrocompound-associated oxidoreductases, NAD(P)H-dependent flavin oxidoreductases or related redox proteins were retrieved as an initial candidate set. Candidate sequences were examined using NCBI Conserved Domain Search to confirm correspondence with the broader YrpB/NMO-associated protein space, including COG2070/YrpB, cd04730/NPD_like and PF03060/NMO domain models^13–15^. At this stage, conserved-domain information was used only to support candidate inclusion; numerical comparison of domain E-values was performed subsequently.

### Phylogenetic analysis of YrpB/NMO-associated proteins

The resulting candidate proteins were used for phylogenetic analysis. An initial maximum-likelihood analysis was restricted primarily to proteins originating from the *E. fetida*-associated system, together with selected experimentally characterized Class I NMO proteins, to examine sequence relationships within the experimental continuum. A second, expanded analysis incorporated additional YrpB/NMO-associated proteins from taxonomically diverse bacteria, including representatives from *Pseudomonas*, *Psychrobacter*, *Serratia*, *Bacillus* and *Mycobacterium*, to place the earthworm-associated proteins within a broader sequence context^5,22,23^.

Amino-acid sequences were aligned using MUSCLE^16^, and maximum-likelihood phylogenies were reconstructed in MEGA^17^ using the Jones–Taylor–Thornton substitution model with 1,000 bootstrap replicates^18^. The resulting trees were used to examine the distribution of the *E. fetida*-associated proteins relative to experimentally characterized NMO proteins, annotated YrpB/NMO-family proteins and functionally unresolved homologues. Phylogenetic placement was used to assess sequence relationships and was not considered sufficient for biochemical functional assignment.

### Comparative genome-level YrpB/NMO-associated repertoires

Following phylogenetic analysis, the number and distribution of YrpB/NMO-associated candidate proteins were compared among the selected genome-resolved strains. This analysis distinguished genomes containing relatively restricted candidate repertoires from those encoding multiple paralogous or related oxidoreductases. *Serratia* sp. EWG9 and *Streptomyces* sp. EAG2 each contained a comparatively simple candidate repertoire, whereas *Mycobacterium* sp. EPG1 and *Pradoshia eiseniae* EAG3 encoded multiple YrpB/NMO-associated proteins. These genome-level patterns were considered together with the preceding growth phenotypes when selecting out the most suitable strain for more detailed analysis.

### Quantitative conserved-domain comparison of YrpB/NMO-associated oxidoreductases

After establishing the phylogenetic relationships, conserved-domain correspondence was examined quantitatively for representative proteins from the genome-resolved isolates, earthworm-associated metagenomic and MAG-derived datasets and selected reference proteins. Each sequence was analysed using NCBI CD-Search^13–15^, and the E-values corresponding to COG2070/YrpB, cd04730/NPD_like and PF03060/NMO were recorded.

For comparative visualization, E-values were transformed as −log_10_(E-value), such that larger values represented stronger correspondence to the respective domain model. The transformed values were displayed as a heatmap. For each protein, the domain model producing the highest −log_10_(E-value) was designated the dominant CDD profile for comparative purposes only.

For clarity, the single focal YrpB/NMO-associated oxidoreductase from *Serratia* sp. EWG9 included in the comparative phylogenetic analyses was designated OXR01 throughout this study (protein accession MBZ0049149; locus tag ITX55_RS20600).

Experimentally characterized proteins were included as functional reference points, including *Pseudomonas aeruginosa* PA4202, a validated Class I nitronate monooxygenase, and PA1024, an experimentally characterized NADH:quinone reductase representing a non-NMO YrpB-family function. Because experimentally distinct proteins can occupy overlapping YrpB/NMO-associated domain space, neither the dominant CDD profile nor the relative E-value pattern was treated as a biochemical functional assignment.

### Multiple-sequence alignment and motif comparison of EWG9 OXR01

The amino-acid sequence of the focal Serratia sp. EWG9 oxidoreductase OXR01 (MBZ0049149) was compared with selected Class I NMO reference sequences, including *Pseudomonas aeruginosa* NMO (PaNMO; NP_252891.1), *Psychrobacter* sp. ANT206 NMO (PsNMO/rPsNMO)^22^, *Klebsiella pneumoniae* NMO (KpNMO; WP_004179795.1) and the *Cellulophaga* NMO sequence WP_013550013.1. Multiple-sequence alignment was performed using BioEdit v7.0^24^ and visualized with ESPript 3.0, following the sequence-comparison framework used for PsNMO characterization (19, 20).

Conserved residues were displayed according to sequence identity and similarity, and the alignment was examined for correspondence with the four conserved sequence regions reported for Class I nitronate monooxygenases^5,22,25^. Residues corresponding to the reported Class I NMO catalytic/conserved positions, including Met18, Asn67, His176, Tyr297 and Lys305 in the PsNMO reference framework, were mapped onto the alignment for comparative assessment of EWG9 OXR01. The analysis was used to assess sequence-level conservation between EWG9 OXR01 and characterized or annotated NMO proteins and was not used independently to assign nitronate monooxygenase activity.

### Physiological characterization of *Serratia* sp. EWG9 under 3-NPA-associated conditions

Growth of *Serratia* sp. EWG9 was examined under minimal-medium conditions designed to distinguish the nutritional context in which 3-nitropropionic acid (3-NPA) supported bacterial proliferation. The basal minimal salt medium (MSM) contained KH□PO□(3 g l□¹), Na□HPO□(6 g l□¹), NaCl (5 g l□¹) and MgSO□·7H□O (0.1 g l□¹). Glucose (8 g l□¹) was supplied as the carbon source and 3-NPA (0.06 g l□¹) replaced inorganic nitrate for the sole-nitrogen-source condition^26,27^. For the sole-carbon-source condition, 3-NPA (0.09 g l□¹) was supplied as the carbon source in the presence of KNO□ (0.875 g l□¹) as the nitrogen source, whereas for the sole-carbon-and-nitrogen-source condition, both glucose and KNO□ were omitted and 3-NPA (0.12 g l□¹) constituted the principal supplied carbon- and nitrogen-containing substrate. Growth was followed for 48 h by viable-cell enumeration and expressed as log□□ CFU ml□¹. *Escherichia coli* K-12 was examined in parallel under the same three nutritional configurations as a comparator. Viable counts were determined over 48 h using the same culture and enumeration procedure, with three independent biological replicates.

To assess concentration-dependent tolerance and growth, EWG9 was additionally cultivated in glucose-containing MSM in which 3-NPA was supplied as the sole nitrogen source at 0.06, 0.3 or 0.6 g l□¹. The concentration series and 48-h growth profiles correspond to Fig. 5b. Cultures were incubated at 30□°C with shaking at 100 rpm. At each sampling point, aliquots were serially diluted in sterile 0.85% NaCl, spread onto Luria agar and incubated at 30□°C before enumeration of colonies. Growth was expressed as log□□ CFU ml□¹. Growth experiments were performed in triplicate.

### Targeted UHPLC–ESI–MS/MS quantification of residual 3-NPA

To determine whether growth of EWG9 was accompanied by depletion of the parent 3-NPA pool, residual 3-NPA in culture supernatants was quantified at 0, 9 and 36 h. An uninoculated abiotic control and *Escherichia coli* K-12 were analysed in parallel to assess non-biological loss and depletion unrelated to the EWG9 phenotype. The analysed profiles correspond to starting concentrations of approximately 59.95 mg l□¹, with EWG9 showing progressive parent-compound loss during incubation.

Quantification was performed using an Acquity reverse-phase UHPLC system coupled to a Waters Xevo tandem-quadrupole mass spectrometer equipped with an electrospray-ionization source. Chromatographic separation was achieved using a Waters BEH C18 column (50 × 2.1 mm, 1.7 µm). Mobile phase A consisted of acetonitrile and mobile phase B consisted of aqueous 5% methanol containing 0.2% formic acid. Separation was performed isocratically at 50:50 (A:B) over 0–3 min at a flow rate of 0.350 ml min□¹. The column and autosampler were maintained at 35□±□1 °C and 20□±□2 °C, respectively, and the injection volume was 2 µl.

3-NPA was detected in multiple-reaction-monitoring mode using a cone voltage of 25 V and collision energy of 8 V. Quantification was based on external calibration with 1/x-weighted linear regression. The final calibration relationship was y=306.36x+809.88, where y represents peak area and x represents 3-NPA concentration. The analytical limits of detection and quantification were 6.76 and 20.47 µg ml□¹, respectively. Samples were diluted in methanol prior to quantification (0.787 ml sample + 0.213 ml MeOH). The analysis was used specifically to quantify disappearance of the parent 3-NPA compound and was not interpreted as evidence of complete mineralization or identification of downstream transformation products.

### Composite RNA-seq analysis of *Serratia* sp. EWG9 under 3-NPA-associated growth

To characterize the early transcriptional response of *Serratia* sp. EWG9 to 3-NPA, cultures were compared under a common glucose background in which either KNO or 3-NPA served as the nitrogen source. A purified colony of EWG9 was initially grown overnight in 10 ml Luria broth at 30□°C and 100 rpm. An aliquot of the overnight culture was washed twice with sterile 0.85% NaCl and transferred to 50 ml minimal salt medium (MSM) containing glucose (4 g l□¹) and KNO (0.875 g l□¹) for 12 h. Cells were then washed twice with sterile 0.85% NaCl and inoculated into either glucose–KNO MSM, containing glucose (8 g l□¹) and KNO (0.875 g l□¹), or glucose–3-NPA MSM, containing glucose (8 g l□¹) and 3-NPA (0.06 g l□¹). Cultures were incubated at 30□°C and 100 rpm in an orbital shaker.

Three independent cultures were established for each condition. Because EWG9 entered logarithmic growth at different times under the two nitrogen regimes, cultures were harvested at their respective early logarithmic-growth phases: 3 h for the glucose–KNO control and 6 h for the glucose–3-NPA condition (A parallel short-interval growth experiment, monitored at 2-h intervals over 24 h under the same glucose–3-NPA condition, was used to provide physiological context for the 6-h transcriptome sampling point). Equal volumes from the three independently grown cultures within each condition were pooled before RNA extraction, generating one composite glucose–KNO□ sample and one composite glucose–3-NPA sample. Cells were collected by centrifugation at 1,500 × *g* for 10 min, suspended in TRIzol and stored at −20□°C before RNA processing^28,29^.

Total RNA was extracted from the composite samples using a TRIzol-based procedure. RNA quality was assessed by denaturing agarose-gel electrophoresis and concentration was determined spectrophotometrically using a SPECTROstar Nano instrument (BMG Labtech, Germany). Messenger RNA was enriched using the MICROBExpress workflow, and sequencing libraries were prepared using the Illumina TruSeq mRNA Sample Preparation workflow. First-strand cDNA synthesis was performed using random hexamer primers and SuperScript III reverse transcriptase, followed by second-strand synthesis. Double-stranded cDNA was purified using AMPure XP beads and fragmented using a Covaris M220 system to an average fragment size of approximately 350 bp. Fragmented cDNA underwent end repair, A-tailing, adapter ligation and limited-cycle PCR amplification before paired-end Illumina sequencing (2 × 150 bp)^28^.

Raw sequencing reads were processed using Trimmomatic v0.38 to remove adapter sequences and low-quality reads. Quality-filtered reads were mapped to the previously reported whole-genome sequence of *Serratia* sp. EWG9 (DDBJ/ENA/GenBank accession JADNRO000000000) using TopHat v2.1.1 with default parameters. Transcript assembly and expression quantification were performed using Cufflinks v2.2.1, and condition-level expression differences between glucose–3-NPA and glucose–KNO□ were calculated using Cuffdiff v2.2.1. Functional assignments were examined using the KEGG Automatic Annotation Server (KAAS), together with manual interrogation of genes associated with replication and translation, glycine/C1 metabolism, respiratory and redox functions, nitrate/nitrite-associated nitrogen handling, carbon and energetic metabolism, and the focal NMO-like YrpB-family oxidoreductase, OXR01 (locus tag ITX55_RS20600).

Because biological cultures were pooled before RNA extraction, the resulting RNA-seq dataset represents one composite expression profile per condition rather than replicate-resolved transcriptomes. Expression changes were therefore used as a condition-level response map to identify coordinated transcriptional patterns associated with early-logarithmic growth under 3-NPA. Cuffdiff-derived fold changes and adjusted *q* values were retained as descriptive outputs of the analytical pipeline, but gene-level statistical significance was interpreted cautiously because biological variance cannot be estimated independently after pooling. In addition, because the □±□ and 3-NPA cultures were sampled at 3 h and 6 h, respectively, the comparison represents transcriptional states at their respective early logarithmic-growth phases rather than a chronologically time-matched experiment.

The integrated response model was constructed from selected condition-associated transcriptional changes together with the physiological growth and parent 3-NPA depletion data. Genes were grouped into functional modules representing replication/translation, glycine/C1 metabolism, respiratory/redox response, nitrate/nitrite-associated functions, and carbon/energetic redistribution. The focal oxidoreductase OXR01 was included in the model as a separate candidate locus for comparison with the broader transcriptional response.

### Molecular docking and protein–ligand interaction analysis

Comparative molecular docking was performed to evaluate the predicted interaction of 3-nitropropionic acid (3-NPA) and related nitro-containing compounds with NMO/YrpB-family oxidoreductases from *Serratia* sp. EWG9, *Pseudomonas aeruginosa* PAO1, *P. aeruginosa* CD3 and *Bacillus subtilis* MTCC121. The ligand panel comprised 3-NPA, nitromethane, nitroethane, 1-nitropropane, 2-nitropropane, chloropicrin, tetranitromethane, 3-nitropropanol, miserotoxin, 2-nitro-2-butene, 2,4,6-trinitrotoluene (TNT), tetryl, nitroglycerin and RDX.

Protein structures were prepared as receptor models using AutoDock Tools v1.5.7^30^. Water molecules were removed, polar hydrogen atoms were added and Kollman charges were assigned. Ligands were energy-minimized and converted to PDBQT format. The three-dimensional structure of 3-NPA was retrieved from PubChem (CID 1678) before preparation for docking^31^.

Because experimentally defined substrate-binding sites were not available for all four proteins, blind docking was performed using AutoDock Vina^32^. A search space of 40 × 40 × 40 Å³ was centred individually on each receptor to encompass the protein surface and potential ligand-accessible regions. Docked conformations were ranked according to the AutoDock Vina scoring function, expressed in kcal mol ¹, and the highest-ranked pose for each protein–ligand combination was retained for comparative analysis. Vina scores were interpreted as computational ranking metrics rather than experimentally determined binding free energies. The broader nitro-compound panel was used only for comparative docking, whereas subsequent interaction profiling and molecular-dynamics analyses were restricted to the 3-NPA complexes.

The highest-ranked EWG9 YrpB–3-NPA, PAO1 NMO–3-NPA, CD3 NMO–3-NPA and *B. subtilis* YrpB–3-NPA complexes were analysed using the Protein–Ligand Interaction Profiler (PLIP)^33^. Predicted hydrogen bonds and other non-covalent contacts were identified from the PLIP interaction profiles and inspected in PyMOL^34,35^. Participating residues, interaction geometry and hydrogen-bond number were compared among the four complexes. Hydrogen-bond counts were treated as descriptors of the predicted interaction network and not as quantitative measures of binding affinity or catalytic activity.

### Molecular-dynamics simulations of the 3-NPA complexes

The highest-ranked 3-NPA docking pose for each protein was subjected to molecular-dynamics (MD) simulation using GROMACS^36,37^ through the Galaxy Europe platform (Galaxy tool version 2022+galaxy0)^11^. Protein parameters were described using the AMBER99SB force field. The 3-NPA ligand was parameterized using the General AMBER Force Field (GAFF), with partial atomic charges assigned using the AM1-BCC charge model^11^. Each protein–3-NPA complex was solvated in a triclinic box containing TIP3P water molecules and electrically neutralized by addition of the required counterions.

Each solvated system was energy-minimized using the steepest-descent algorithm for a maximum of 50,000 steps, with an energy-minimization tolerance of 1,000 kJ mol ¹ nm ¹ and a maximum minimization step size of 0.01 nm. Long-range electrostatic interactions were treated using smooth Particle-Mesh Ewald electrostatics, with short-range Coulomb, neighbour-list and van der Waals cut-offs of 1.0 nm.

MD simulations were performed under an isothermal–isochoric (NVT) ensemble at 300 K. Initial velocities were assigned at the start of the simulation, and protein and non-protein atoms were treated as separate temperature-coupling groups. Newtonian equations of motion were integrated using the leap-frog algorithm with a timestep of 0.002 ps (2 fs), and bonds involving hydrogen atoms were constrained. Neighbour searching was performed using a buffered pair-list scheme, and long-range electrostatic interactions were treated using fast smooth Particle-Mesh Ewald electrostatics. Coulomb, neighbour-list and short-range van der Waals cut-offs were each maintained at 1.0 nm.

Each protein–3-NPA system was propagated for 150,000,000 integration steps, corresponding to a total simulation time of 300 ns. Simulation data were sampled every 1,000 integration steps, corresponding to a 2-ps interval. Identical simulation settings were applied to the EWG9 YrpB–3-NPA, PAO1 NMO–3-NPA, CD3 NMO–3-NPA and *B. subtilis* YrpB–3-NPA systems to permit direct comparison of their trajectory behaviour.

### Analysis of the 300-ns MD trajectories

Global conformational behaviour was evaluated using backbone root-mean-square deviation (RMSD) over the complete 300-ns trajectories. Trajectory frames were structurally fitted to the reference protein structure before RMSD calculation to remove overall translational and rotational motion, and RMSD values were expressed in nm. In addition to whole-trajectory assessment, mean RMSD values were summarized over 0–100, 100–200 and 200–300 ns intervals to distinguish early, intermediate and late conformational behaviour.

Residue-level flexibility was assessed using root-mean-square fluctuation (RMSF) after removal of overall translational and rotational motion. RMSF values were expressed in nm and used to identify localized regions of increased or reduced conformational mobility.

Overall structural compactness was evaluated from the radius of gyration (Rg) throughout each trajectory. Rg values were expressed in nm and used to assess whether trajectory-dependent conformational changes were accompanied by global expansion or contraction of the protein structure.

Potential-energy trajectories were extracted from the GROMACS simulations^36,37^ and expressed in kJ mol ¹. Potential energy was evaluated primarily for temporal stability and sustained energetic drift. Because independently solvated systems differed in composition and total atom number, absolute potential-energy values were not interpreted as measures of relative protein–ligand binding strength.

For comparative visualization of energetic fluctuations, each potential-energy trajectory was mean-centred by subtracting its trajectory-wide mean. This transformation was used solely to facilitate comparison of temporal fluctuations among systems; all quantitative analyses were based on the original, untransformed energy values.

### MD-derived binding-pocket analysis using fpocket online server

Binding-pocket architecture was evaluated from the MD-derived protein conformations using fpocket online server^38,39^. For each protein–3-NPA system, the final structure at 300 ns was extracted and subjected to pocket detection. The highest-ranked predicted cavity, designated Pocket 1, was retained for comparative analysis.

Pocket descriptors included fpocket score, druggability score, cavity volume, number of alpha spheres, total solvent-accessible surface area (SASA), polar and apolar SASA, proportion of polar atoms, polarity score, charge score, hydrophobicity-related descriptors and mean local hydrophobic density. For example, the EWG9 Pocket 1 output comprised a volume of 960.360 Å³, 153 alpha spheres and a druggability score of 0.913, confirming that these quantities were taken directly from the fpocket output. Corresponding Pocket 1 descriptors were obtained independently for the comparator proteins; PAO1, for example, yielded a volume of 1,642.163 Å³ and 216 alpha spheres.

For comparative visualization of Pocket 1 architecture, cavity volume, number of alpha spheres and mean local hydrophobic density were displayed directly from the fpocket output.

No additional normalization was applied to the other displayed pocket descriptors. fpocket and druggability scores were interpreted as structural ligandability descriptors rather than estimates of 3-NPA binding free energy or catalytic activity. Pocket structures and alpha-sphere distributions were visualized in PyMOL for comparison among EWG9 YrpB, PAO1 NMO, CD3 NMO and *B. subtilis* YrpB.

### Statistical analysis

Statistical analyses were performed in IBM SPSS Statistics v27^40,41^ using log_10_ CFU ml^−1^ values from three biological replicates. Nutritional-condition time-course data were analysed using linear mixed-effects models with condition, centred time, quadratic time and their interactions as fixed effects and biological culture as a random intercept. Balanced concentration- and strain-dependent time courses were analysed by two-way mixed-design repeated-measures ANOVA, with time as the within-subject factor and concentration or strain as the between-subject factor. Sphericity was assessed using Mauchly’s test, with Greenhouse–Geisser correction where required, and Bonferroni adjustment was applied to pairwise comparisons. The 2-h interval time course was analysed using linear and quadratic time effects with a first-order autoregressive [AR(1)] covariance structure for repeated observations within cultures. For *E. coli* K-12, comparisons among nutritional conditions were based on the common 0, 24 and 48 h sampling points. P < 0.05 was considered significant.

## RESULTS

### YrpB-associated oxidoreductases dominate a heterogeneous environmental YrpB/NMO sequence space

Across 13 metagenomes representing earthworm/compost-, host-, soil- and waste-associated environments, 390 conserved-domain-supported YrpB/NMO-associated proteins were recovered (Fig. 1a,b, Supplementary Table 2). Of these, 324 (83.1%) were classified operationally as YrpB-associated and 66 (16.9%) as NMO-like. YrpB-associated proteins were detected in all datasets, whereas NMO-like proteins were absent from three. This predominance was retained after normalization to the number of annotated proteins: YrpB-associated abundance exceeded NMO-like abundance in every metagenome and ranged from 0.93 to 4.59 hits per 10,000 annotated proteins. Within individual datasets, YrpB-associated proteins comprised 64.3–100% of the combined YrpB/NMO candidate pool. Thus, the conserved-domain-supported environmental YrpB/NMO-associated candidate pool was consistently YrpB-dominated rather than NMO-dominated.

**Fig. 1:**
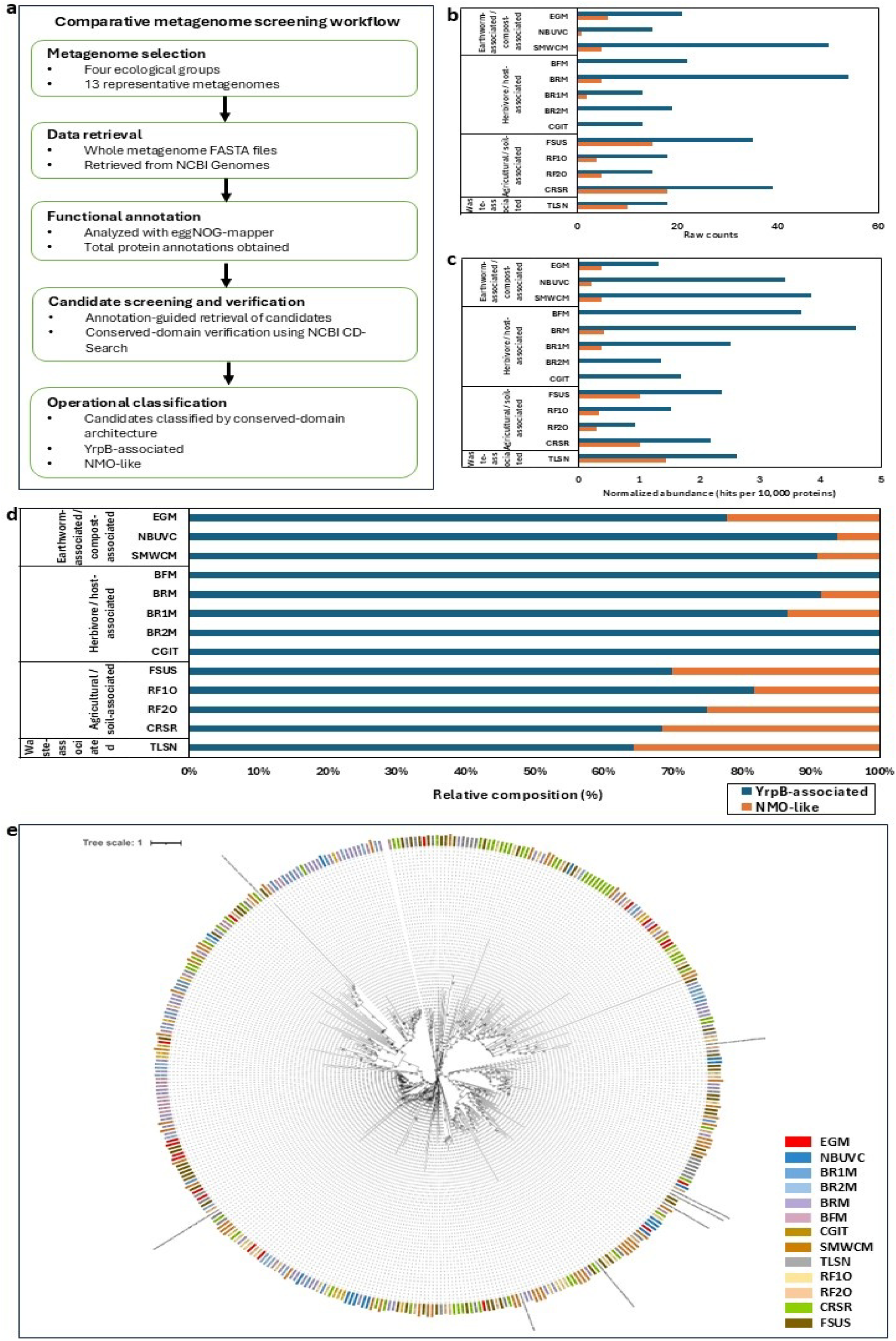
Environmental distribution and phylogenetic diversity of YrpB/NMO-associated flavin oxidoreductases across representative metagenomes. **a,** Workflow used for comparative screening of 13 independently sourced metagenomes representing earthworm- and compost-associated, herbivore/host-associated, agricultural/soil-associated and waste-associated environments. Whole-metagenome protein sets were annotated using eggNOG-mapper, candidate nitrocompound-associated oxidoreductases were retrieved by conserved-domain-guided screening. Candidates were operationally grouped as YrpB-associated or NMO-like according to annotation and conserved-domain correspondence; these categories are used for comparative screening and do not by themselves establish biochemical function. **b,** Raw numbers of YrpB-associated and NMO-like candidate proteins recovered from each metagenome. **c,** Candidate abundance normalized to the total number of annotated proteins in each dataset and expressed as hits per 10,000 proteins. **d,** Relative composition of the two operational candidate groups within each metagenome, calculated as the proportion of YrpB-associated or NMO-like candidates among their combined total. **e,** Maximum-likelihood phylogeny of the YrpB/NMO-associated proteins recovered from the 13 metagenomes together with selected reference proteins included to provide functional and phylogenetic context. Metagenome-derived sequences are indicated by source-specific-coloured labels, whereas reference proteins are shown in **bold**. Reference sequences include experimentally validated Class I nitronate monooxygenases, functionally unresolved NMO/YrpB-COG2070 homologues, YrpB-like COG2070-family oxidoreductases. The tree illustrates extensive sequence diversification across environmental and host-associated sources and places the environmental candidates relative to experimentally characterized and functionally unresolved members of the broader YrpB/COG2070–NMO sequence space. Ecological groupings are descriptive, and individual metagenomes represent independent datasets rather than biological replicates. EGM, earthworm gut metagenome; NBUVC, North Bengal University vermicompost; SMWCM, soil amended with raw cow manure; BFM, bovine faecal metagenome; BRM, bovine rumen metagenome; BR1M and BR2M, buffalo rumen metagenomes 1 and 2; CGIT, chicken gastrointestinal tract metagenome; FSUS, farmland soil, USA; RF1O and RF2O, rice-field soil metagenomes 1 and 2, Odisha; CRSR, cotton rhizosphere soil, Rajasthan; TLSN, top landfill soil, Nagpur.

Maximum-likelihood analysis further distributed the 390 environmental proteins across multiple branches of the broader YrpB/COG2070–NMO sequence space rather than a single compact lineage (Fig. 1e). Environmental candidates occurred within a sequence space containing both the experimentally validated Class I NMO PA4202 and the non-NMO YrpB-family NADH:quinone reductase PA1024, together with numerous unresolved homologues. This phylogenetic heterogeneity indicated that annotation within this protein family alone does not resolve biochemical function.

To examine this diversity within a tractable soil-associated system, we next focused on an *Eisenia fetida* feed–gut–cast continuum. Earthworms occupy the soil–detritus interface, substantially restructure associated microbial communities and are widely used in cattle-manure-based vermicomposting. Their feed–gut–cast progression can also be reproduced experimentally, providing a framework in which bacterial transitions and associated protein repertoires can be followed across successive ecological compartments.

### Cultivable 3-NPA-responsive bacteria span diverse lineages across the feed–gut–cast continuum

Selective cultivation recovered 31 reproducibly 3-NPA-responsive isolates from formulated feed, whole gut, freshly produced casts and the corresponding normal-feed continuum (Fig. 2a and Supplementary Table 3). Nearly full-length 16S rRNA gene analysis showed substantial taxonomic diversity among these isolates. For comparison with the culture-independent community, the present collection was combined with previously recovered legacy isolates, yielding a broader cultivable inventory spanning 24 genera. Thus, growth under the 3-NPA-containing condition was not restricted to a narrow taxonomic lineage. Isolates from the formulated-feed, gut, cast and control compartments were distributed across multiple branches of the 16S maximum-likelihood tree.

**Fig. 2:**
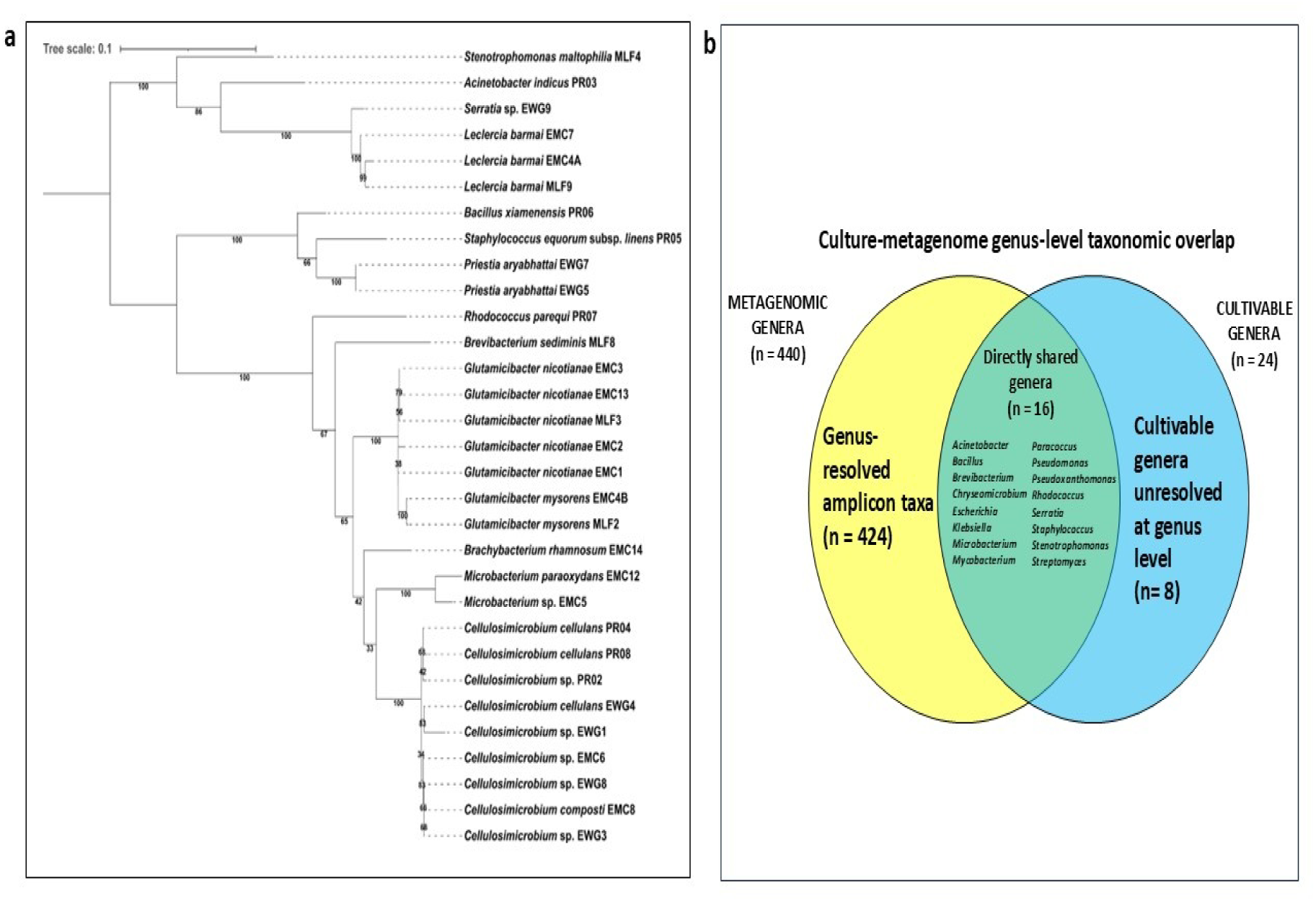
Taxonomic diversity and culture–amplicon correspondence of cultivable 3-NPA-responsive bacteria from the *Eisenia fetida* feed–gut–cast continuum. **a,** Maximum-likelihood phylogeny of 31 cultivable 3-NPA-responsive bacterial isolates based on Sanger-sequenced 16S rRNA gene amplicons. The tree was reconstructed in MEGA12 with 1,000 bootstrap replicates; bootstrap support values are shown at internal nodes and the scale bar indicates substitutions per site. Isolate prefixes denote their source within the feed–gut–cast systems: MLF, formulated feed; EWG, whole gut of formulated-feed-fed *E. fetida*; EMC, casts from formulated-feed-fed worms; and PR, isolates from the normal-feed control continuum. **b,** Genus-level correspondence between the cultivable collection and the culture-independent V3-region 16S rRNA gene amplicon inventory of the corresponding feed–gut–cast system. The amplicon dataset contained 440 genera resolved at genus level, whereas the cultivable collection represented 24 genera. Sixteen cultivable genera were directly represented in the amplicon-derived genus inventory, leaving 424 other genus-resolved amplicon taxa. Eight cultivable genera were not resolved to genus level in the V3 amplicon dataset; corresponding higher-rank family- and/or class-level assignments were nevertheless represented and these taxa were therefore treated as unresolved at genus level rather than demonstrably absent from the culture-independent community.

The accompanying *in silico Hae*III restriction profiles showed marked variation among isolates and broadly reflected the sequence diversity represented in the 16S tree (Fig. 2a and Supplementary Fig. 2). Because these profiles were computationally derived from the same amplicon sequences, they were interpreted as a complementary visualization of sequence heterogeneity rather than independent phylogenetic evidence.

Comparison with culture-independent V3-region 16S datasets further showed that the cultivable collection recovered a substantial subset of the feed–gut–cast community (Fig. 2b and Supplementary Table 4). Sixteen of the 24 cultivable genera (66.7%) were also detected at genus level among the 440 genus-resolved taxa in the amplicon dataset, including *Acinetobacter, Bacillus, Brevibacterium, Chryseomicrobium, Escherichia, Klebsiella, Microbacterium, Mycobacterium, Paracoccus, Pseudomonas, Pseudoxanthomonas, Rhodococcus, Serratia, Staphylococcus, Stenotrophomonas* and *Streptomyces*. The remaining cultivable genera were represented at higher taxonomic ranks but were not resolved to genus level in the V3 dataset. The cultivable 3-NPA-responsive assemblage therefore represented multiple lineages embedded within the broader feed–gut–cast community.

### EWG9 combines rapid generation with sustained growth under 3-NPA-associated conditions

Generation-time screening of the 31 isolates revealed substantial phenotypic variation under glucose– 3-NPA conditions, with mean generation times ranging from approximately 3.08 to 7.67 h (Fig. 3a). *Serratia* sp. EWG9 showed one of the shortest generation times (3.079 h), essentially matching EMC7 (3.081 h), and was therefore selected as one of the fastest 3-NPA-responsive isolates. EWG9 retained strongest growth around neutral-to-mildly alkaline pH and at 30–37 °C (Supplementary Fig. 3), when evaluated.

**Fig. 3:**
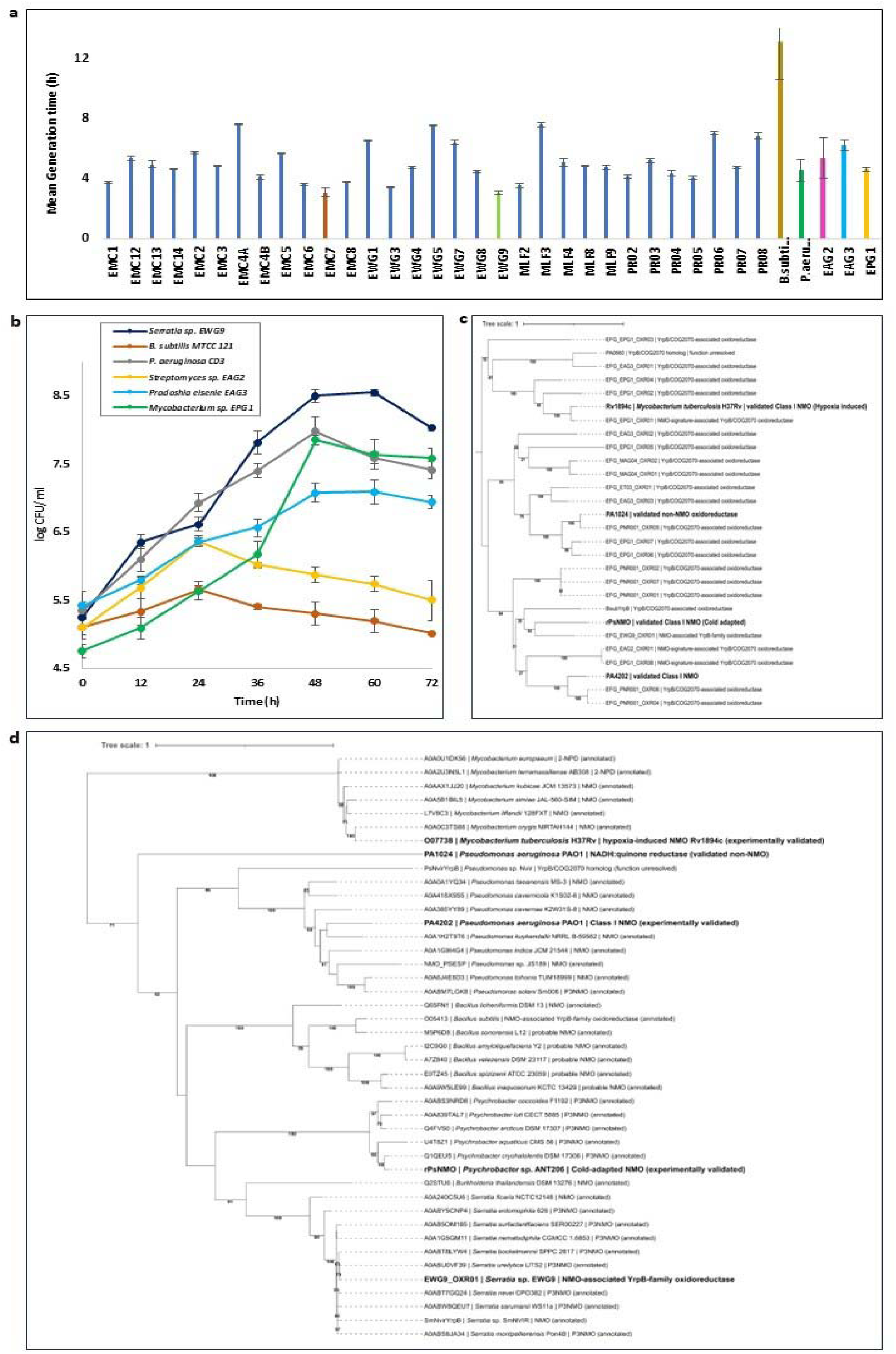
Phenotypic variation among cultivable 3-NPA-responsive bacteria is accompanied by phylogenetic diversity of YrpB/NMO-associated oxidoreductases. **a,** Mean generation times of 31 cultivable 3-NPA-responsive isolates recovered from the *Eisenia fetida* feed–gut–cast continuum, together with selected comparator strains. Bars show mean generation time and error bars indicate SD among replicate measurements. *Serratia* sp. EWG9 was among the fastest-growing isolates under the screening condition, while generation times varied substantially across the cultivable collection. **b,** Comparative viable-count profiles of selected genome-resolved 3-NPA-responsive isolates and comparator strains during growth in glucose–3-NPA mineral salts medium. *Serratia* sp. EWG9 showed sustained population increase over 72 h relative to *Mycobacterium* sp. EPG1, *Pradoshia eiseniae* EAG3, *Streptomyces* sp. EAG2, *Pseudomonas aeruginosa* CD3 and *Bacillus subtilis* MTCC121. Data are mean ± SD (*n* = 3 independent biological replicates). Two-way mixed-design repeated-measures ANOVA showed significant effects of time, strain and time × strain (*F□□*,□□= 84.67, *P* < 0.001); Bonferroni-adjusted comparisons showed that EWG9 had a higher overall viable count than all comparator strains. **c,** Maximum-likelihood phylogeny of YrpB/COG2070–NMO-associated proteins from selected genome-resolved isolates together with functionally informative reference proteins. Experimentally validated Class I NMOs, characterized non-NMO COG2070 oxidoreductases and functionally unresolved homologues are included to provide sequence context for the oxidoreductase repertoires of the cultivable strains. **d,** Expanded maximum-likelihood phylogeny placing EWG9 OXR01 and selected YrpB/NMO-associated proteins within a broader set of bacterial homologues from *Mycobacterium*, *Pseudomonas*, *Bacillus*, *Psychrobacter*, *Burkholderia* and *Serratia*. EWG9 OXR01 occurs within a *Serratia*-enriched NMO-associated lineage containing proteins predominantly annotated as nitronate monooxygenases or propionate-3-nitronate monooxygenases, whereas experimentally validated Class I NMOs occupy distinct taxon-associated lineages. Functionally characterized reference proteins are highlighted in bold. For c and d, bootstrap support values are shown at internal nodes and scale bars indicate amino-acid substitutions per site.

A focused 72-h comparison with genome-resolved earthworm-associated isolates and additional NMO/YrpB-background comparators further distinguished EWG9 (Fig. 3b). EWG9 increased from ∼5.26 log_10_CFU ml^−1^ at inoculation to 7.82 by 36 h, reached ∼8.50–8.55 log CFU ml^−1^ at 48–60 h and remained above 8 log CFU ml^−1^ at 72 h. *Pseudomonas aeruginosa* CD3 and *Mycobacterium* sp. EPG1 also showed substantial growth but reached lower maxima of ∼7.98 and ∼7.85 log CFU ml^−1^, respectively. *Pradoshia eiseniae* EAG3 increased more gradually, *Streptomyces* sp. EAG2 showed a lower early maximum followed by decline, and *Bacillus subtilis* MTCC121 displayed comparatively limited net expansion. EWG9 therefore combined rapid generation with sustained population growth, providing a phenotypic basis for deeper genome- and protein-level analysis.

### EWG9 OXR01 occupies a heterogeneous YrpB/NMO-associated protein space

Phylogenetic analysis of YrpB/NMO-associated proteins from the *E. fetida*-linked system revealed multiple distinct sequence lineages rather than a unified NMO-associated clade (Fig. 3c,d). The multiple EPG1 proteins were distributed across several branches, including lineages near experimentally validated Class I NMOs, while the three EAG3 proteins were also separated across the tree, indicating within-genome sequence diversification.

EWG9 encoded a single focal candidate, OXR01, which clustered within a lineage containing the experimentally characterized PsNMO and a *Bacillus* YrpB-family homologue (Fig. 3c). In an expanded phylogeny incorporating broader taxonomic representation, EWG9 OXR01 instead grouped within a well-supported *Serratia*-dominated lineage containing proteins annotated as NMO or P3NMO (Fig. 3d). Experimentally validated NMOs from *Pseudomonas, Psychrobacter* and *Mycobacterium* occupied separate taxon-associated branches. Thus, EWG9 OXR01 clearly belonged to the wider YrpB/NMO-associated sequence family, but its position did not establish biochemical equivalence to a validated NMO. This ambiguity motivated the subsequent genome-level repertoire and conserved-domain comparisons, used to determine how EWG9 differed from the other phenotypically selected strains (Fig. 4a and Supplementary Table 5).

**Fig. 4:**
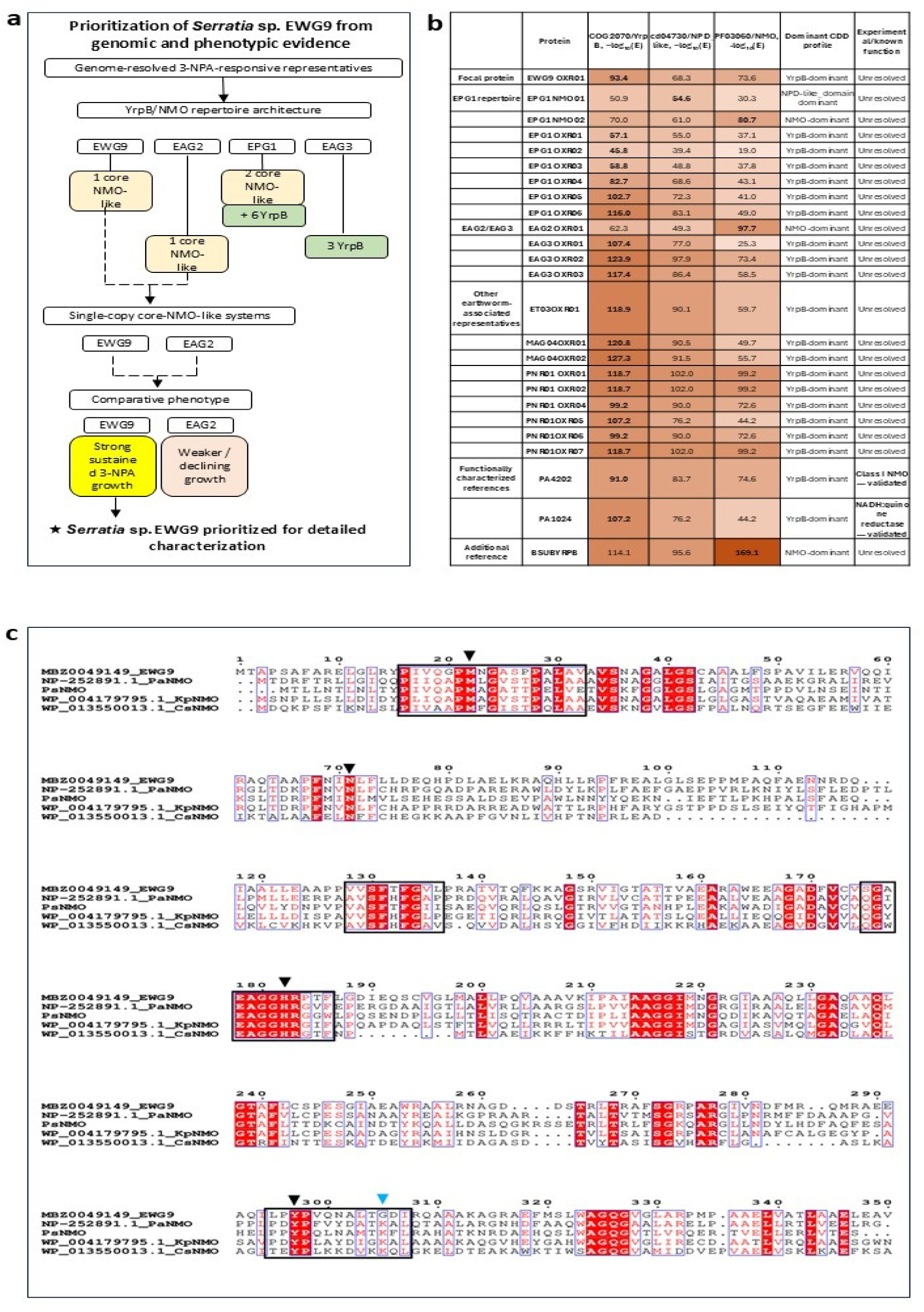
Prioritization of *Serratia* sp. EWG9 and sequence-level evaluation of its candidate YrpB/NMO-family oxidoreductase. **a,** Decision framework used to prioritize genome-resolved 3-NPA-responsive representatives for detailed study. Comparative genome screening showed that EWG9 and EAG2 each encoded a single candidate core NMO-like/YrpB–COG2070 oxidoreductase, whereas EPG1 carried a larger mixed repertoire and EAG3 carried only broader YrpB-like representatives. Phenotypic comparison under 3-NPA-associated growth conditions further distinguished EWG9, which showed strong sustained growth, from EAG2, which showed weaker or declining growth. Together, low candidate-copy number and superior phenotype led to prioritization of EWG9 for detailed characterization. **b,** Comparative conserved-domain profile of the focal EWG9 protein, candidate oxidoreductases from representative cultivable isolates and selected reference proteins. Values represent -log (E) values for the best NCBI CDD correspondence to COG2070/YrpB, cd04730/NPD_like and PF03060/NMO models, and the dominant CDD profile was assigned from the strongest correspondence. The comparison includes the focal EWG9 OXR01 protein, the EPG1 repertoire, EAG2/EAG3 representatives, additional earthworm-associated homologs and functionally characterized references. The table shows that experimentally validated proteins can also reside within strongly overlapping YrpB/COG2070-, NPD_like- and NMO-associated domain space, indicating that conserved-domain correspondence alone is insufficient to establish biochemical function. **c,** Multiple-sequence alignment of EWG9 OXR01 with validated and reference Class I NMO-related proteins, including PaNMO, PsNMO, KpNMO and CsNMO. Black boxes denote the four reported conserved Class I NMO motif regions. Black triangles mark the positions corresponding to Met18, Asn67, His176 and Tyr297, which are conserved in EWG9. The blue triangle marks the position corresponding to Lys305 in PsNMO/related Class I NMO references; this position is occupied by Gly in EWG9. Thus, EWG9 retains the overall Class I NMO motif architecture and four key active-site-associated residues, but carries a notable divergence at the Lys305-equivalent position. Together, the architecture, alignment and phenotype support EWG9 OXR01 as a divergent, putative Class I NMO-like YrpB/COG2070 homolog, while its direct catalytic role remains unresolved.

### Conserved-domain and motif analyses retain NMO association but not functional resolution

Quantitative conserved-domain comparison further exposed overlap within the family (Fig. 4b). EWG9 OXR01 showed strong correspondence to all three recurrent domain models, with −log10(E) values of 93.4 for COG2070/YrpB, 68.3 for NPD_like/cd04730 and 73.6 for PF03060/NMO. Although COG2070/YrpB provided the strongest match, the substantial NPD_like and PF03060 signals indicated an overlapping YrpB/NMO architecture.

This ambiguity was also evident among experimentally characterized references. The validated Class I NMO PA4202 itself showed a YrpB-dominant profile (91.0, 83.7 and 74.6 for YrpB, NPD_like and NMO, respectively), whereas the experimentally characterized non-NMO YrpB-family NADH:quinone reductase PA1024 was also YrpB-dominant but showed weaker PF03060 correspondence. EAG2 OXR01 was strongly NMO-dominant, whereas EAG3 proteins were YrpB-dominant and EPG1 contained a mixed repertoire. A dominant YrpB signature was therefore not diagnostic of either NMO or non-NMO activity.

Multiple-sequence alignment added a further sequence-level layer (Fig. 4c). EWG9 OXR01 retained four strongly conserved regions shared with selected Class I NMO references and preserved positions corresponding to several reported NMO-associated residues, including Met18, Asn67, His176 and Tyr297, while showing local divergence around the Lys305-equivalent region. EWG9 OXR01 therefore retained substantial Class I NMO-associated sequence architecture but remained distinct from characterized reference enzymes.

### EWG9 grows across distinct nutritional roles of 3-NPA and depletes the parent compound

EWG9 proliferated under all three tested nutritional configurations of 3-NPA, although growth was strongly context dependent with the strongest increase observed when 3-NPA replaced nitrate as the sole nitrogen-containing substrate. By contrast, *E. coli* K-12 showed declining viable counts under all three corresponding conditions, with the greatest decline when 3-NPA served as the combined carbon and nitrogen source (Fig. 5a and Supplementary Fig. 4). EWG9 additionally maintained robust growth across 0.06, 0.3 and 0.6 g l^−1^ 3-NPA when the compound served as the nitrogen source (Fig. 5b). Because the starting viable counts differed among treatments, these data were not interpreted as a dose-dependent growth advantage, but they demonstrated sustained proliferation across a tenfold concentration range.

**Fig. 5:**
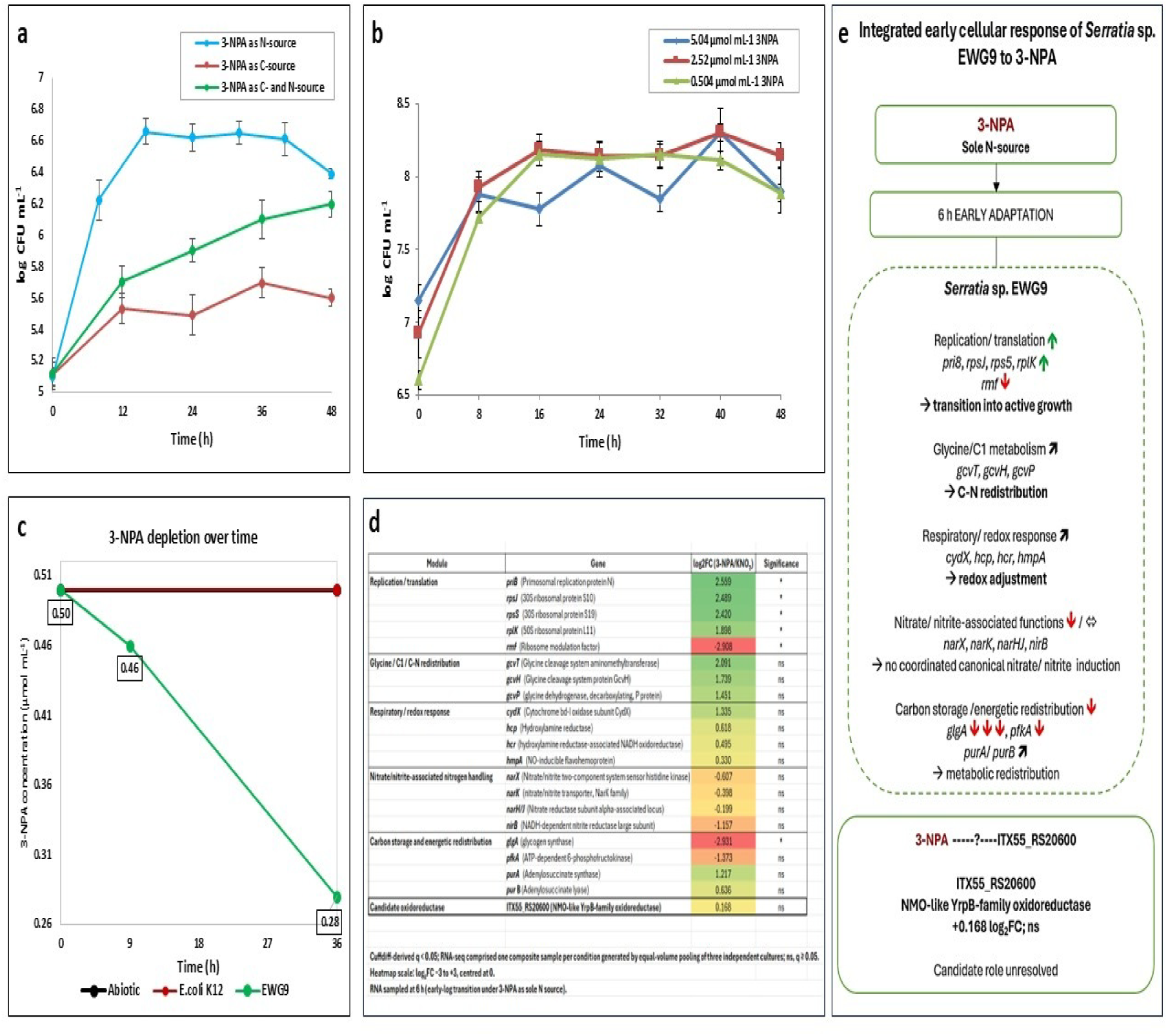
*Serratia* sp. EWG9 couples 3-NPA-supported growth with parent-compound depletion and a distributed early cellular response. **a,** Growth of *Serratia* sp. EWG9 in mineral salts medium containing 3-NPA as the sole nitrogen source, sole carbon source or combined carbon and nitrogen source, determined by viable-cell enumeration over 48□h. Data are mean ± SD (*n* = 3 biological replicates). Linear mixed-effects modelling identified significant condition × time (*P* = 0.010) and condition × time² (*P* < 0.001) interactions, indicating that growth trajectories differed among the three nutritional configurations. **b,** Growth of EWG9 with 3-NPA supplied at 0.504, 2.52 or 5.04□µmol□ml□¹ as the sole nitrogen source in glucose-containing mineral salts medium. Data are mean ± SD (*n* = 3 biological replicates). Two-way mixed-design repeated-measures ANOVA showed significant effects of time (*F*□,□□= 156.40, *P* < 0.001, ηp² = 0.963), 3-NPA concentration (*F*□,□= 27.83, *P* = 0.001, ηp² = 0.903) and time × concentration (*F*□□,□□= 7.08, *P* < 0.001, ηp² = 0.702). Bonferroni-adjusted comparisons showed higher overall viable counts at 2.52□µmol□ml□¹ than at 0.504□µmol□ml□¹ (*P* = 0.003) or 5.04□µmol□ml□¹ (*P* = 0.001). **c,** Depletion of parent 3-NPA by EWG9 compared with *Escherichia coli* K-12 and abiotic controls. Residual 3-NPA decreased from approximately 0.50□µmol□ml□¹ at 0□h to 0.46□µmol□ml□¹ at 9□h and 0.28□µmol□ml□¹ at 36□h in EWG9 cultures, whereas both controls remained close to the starting concentration. Parent 3-NPA was quantified by targeted UHPLC–MS/MS; these measurements demonstrate disappearance of the parent compound but do not identify transformation products or establish complete mineralization. **d,** Selected transcriptional changes during early-logarithmic growth of EWG9 under 3-NPA relative to the KNO□ condition, grouped into replication/translation, glycine/C1 metabolism and C–N redistribution, respiratory/redox response, nitrate/nitrite-associated nitrogen handling, carbon storage and energetic redistribution, and the candidate oxidoreductase OXR01. Values are Cuffdiff-derived log fold changes from one composite RNA-seq sample per condition generated by equal-volume pooling of three independently cultured biological preparations before RNA extraction; asterisks denote *q* < 0.05 and ns denotes *q* ≥ 0.05. The data therefore represent condition-level transcriptional response patterns rather than replicate-resolved differential-expression estimates. **e,** Integrated model of the early cellular response of EWG9 to 3-NPA as the sole nitrogen source. Increased replication/translation-associated expression, shifts in glycine/C1 metabolism, respiratory/redox adjustment and altered carbon-storage and energetic functions occurred without coordinated induction of canonical nitrate/nitrite-associated genes. The NMO-associated YrpB-family oxidoreductase OXR01 showed only a marginal transcriptional change (+0.168 log fold change; ns), leaving its direct contribution to 3-NPA transformation unresolved. The model integrates physiological, parent-compound depletion and composite transcriptomic observations and does not imply catalytic causality for OXR01.

Targeted UHPLC–ESI–MS/MS then showed that this phenotype was accompanied by measurable depletion of parent 3-NPA (Fig. 5c and Supplementary Fig. 5). Residual 3-NPA decreased from ∼0.50 µmol ml^−1^ initially to ∼0.46 µmol ml^−1^ at 9 h and ∼0.28 µmol ml^−1^ at 36 h, corresponding to ∼44% depletion. Abiotic and *E. coli* K-12 controls retained approximately the starting concentration. These measurements establish disappearance of the parent compound, although they do not identify downstream products or demonstrate complete mineralization.

### Early 3-NPA growth is accompanied by a distributed transcriptional response

Composite RNA-seq of EWG9 grown in presence of 3-NPA revealed a distributed early cellular response rather than selective induction of the focal oxidoreductase, while the cells are still adapting to the 3-NPA condition (Fig. 5d,e). Parallel 2-h interval growth profiling showed an early decline in viability at 2 h followed by recovery, contextualizing the 6-h transcriptome as an early post-stress adaptation state (Supplementary Fig. 6). Replication- and translation-associated genes showed the clearest Cuffdiff-supported changes: *priB, rpsJ, rpsS* and *rplK* increased by 2.56, 2.49, 2.42 and 1.90 log2-fold, respectively, whereas *rmf* decreased by 2.91 log2-fold (q<0.05). Glycine/C1-associated genes *gcvT, gcvH* and *gcvP* increased by 2.09, 1.74 and 1.45 log2-fold, although these changes were not significant in the pooled dataset. Respiratory/redox genes, including *cydX, hcp, hcr* and *hmpA*, also showed moderate positive shifts.

By contrast, *narX, narK, narH/J* and *nirB* were not coordinately induced, indicating that replacement of nitrate by 3-NPA did not trigger broad activation of the canonical nitrate/nitrite-associated machinery. Carbon-storage functions also shifted, with *glgA* decreasing by 2.93 log2FC (q<0.05) and *pfkA* decreasing by 1.37 log2FC.

Notably, the focal NMO-like YrpB-family oxidoreductase OXR01 changed only marginally (+0.168 log2FC; ns). Thus, the physiological ability of EWG9 to grow with and deplete 3-NPA was not accompanied by strong induction of this locus at the sampled early-logarithmic stage. Because the transcriptomes were composite samples and harvested at condition-specific early-log phases, these changes are interpreted as condition-level response patterns rather than replicate-resolved differential expression.

### Structural analyses support stable 3-NPA accommodation by EWG9 OXR01

Within the broader ligand panel, EWG9 OXR01 occupied the highest or joint-highest docking rank for the plant-associated nitrotoxins 3-NPA, 3-nitropropanol and miserotoxin, suggesting preferential predicted compatibility with this naturally occurring nitropropionate/nitropropanol chemical space (Supplementary Table 1). For 3-NPA, EWG9 OXR01 produced the most favourable Vina score (−5.3 kcal mol−1), compared with CD3 (−5.1), PAO1 (−5.0) and *B. subtilis* (−4.9). PLIP identified seven predicted hydrogen bonds in the EWG9–3-NPA complex, compared with six for PAO1 and five each for CD3 and *B. subtilis* (Fig. 6a and Supplementary Fig. 1).

**Fig. 6.**
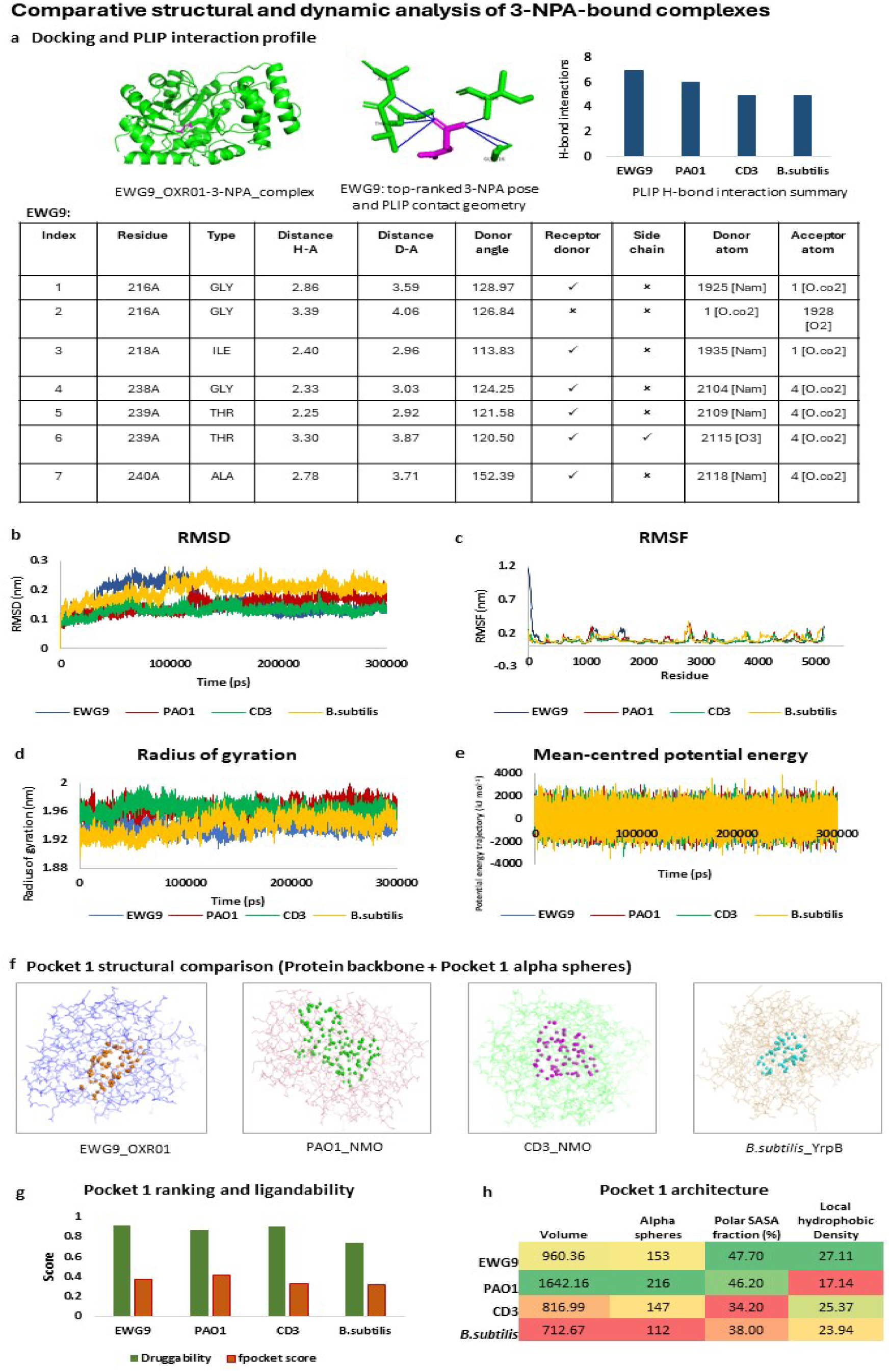
Comparative docking, interaction, molecular-dynamics and pocket analyses support stable 3-NPA accommodation by EWG9 YrpB. **a,** Predicted 3-NPA binding to EWG9YrpB, showing the top-ranked docked complex, representative PLIP contact geometry and hydrogen-bond interactions identified for the EWG9YrpB–3-NPA complex. The accompanying comparison summarizes the number of predicted hydrogen-bond interactions for EWG9YrpB and the corresponding PAO1, CD3 and *Bacillus subtilis* comparator proteins. **b–e,** Comparative 300-ns molecular-dynamics trajectories of the 3-NPA-bound complexes. **b,** Backbone root-mean-square deviation (RMSD), showing early conformational adjustment followed by late-stage stabilization of the four complexes. **c,** Residue-wise root-mean-square fluctuation (RMSF), revealing predominantly localized flexibility superimposed on comparatively restrained global residue fluctuations. **d,** Radius of gyration, showing maintenance of overall protein compactness throughout the simulations. **e,** Potential-energy trajectories, which remained broadly stable during the 300-ns simulations. Together, these parameters indicate that the EWG9 OXR01 protein retained overall conformational stability within the simulated 3-NPA-bound system and compact over the simulation interval, with dynamic behaviour comparable to the reference and comparator systems. **f,** Structural comparison of the highest-ranked Fpocket cavity (Pocket 1) in EWG9YrpB, PAO1 NMO, CD3 NMO and *B. subtilis* YrpB; coloured alpha spheres indicate the predicted pocket volumes within the corresponding protein backbones. **g,** Comparative Fpocket druggability and pocket scores for Pocket 1 across the four proteins. **h,** Pocket 1 architecture summarized by cavity volume, number of alpha spheres, polar solvent-accessible surface-area fraction and local hydrophobic density. EWG9YrpB displayed an intermediate-sized cavity with substantial polar surface contribution and a locally hydrophobic environment compatible with accommodation of a small polar ligand such as 3-NPA. Docking, PLIP, molecular-dynamics and pocket analyses provide structural evidence for ligand compatibility but do not establish catalytic turnover or functional equivalence to a Class I nitronate monooxygenase.

Over 300 ns of molecular-dynamics simulation, all four protein structures retained overall conformational stability during the simulations (Fig. 6b–e). EWG9 showed a progressive reduction in mean RMSD from 0.186 nm during 0–100 ns to 0.130 nm during 200–300 ns, while retaining essentially invariant radius of gyration (∼1.939 nm). Most residues remained comparatively restrained despite localized N-terminal flexibility.

Pocket analysis of the final MD structures identified an intermediate-sized EWG9 cavity (960.36 Å^3^) with the highest druggability score (0.913), high local hydrophobic density and a substantial polar surface contribution (∼47.7%) (Fig. 6f–h). Collectively, docking, residue-level interactions, long-timescale conformational behaviour and cavity architecture support EWG9 OXR01 as a structurally credible 3-NPA-interacting candidate. These computational observations, however, demonstrate ligand compatibility rather than catalytic turnover, leaving the biochemical role of this divergent YrpB/NMO-associated oxidoreductase unresolved.

## DISCUSSION

The present study places bacterial 3-nitropropionic acid (3-NPA) responsiveness within a substantially broader oxidoreductase landscape than would be predicted from a canonical nitronate monooxygenase (NMO)-centred view. Across the environmental metagenomes examined here, YrpB-associated proteins dominated the retrieved YrpB/NMO candidate pool and were dispersed across a heterogeneous phylogenetic space. This pattern should not be interpreted to mean that environmental YrpB homologues are themselves 3-NPA-metabolizing enzymes. Rather, it identifies a large reservoir of related flavin oxidoreductases whose functional boundaries remain incompletely resolved. This distinction is particularly important for NMOs, for which historical sequence-based annotations substantially outpaced biochemical characterization. The structural and kinetic characterization of *Pseudomonas aeruginosa* PA4202 established the first experimentally validated bacterial Class I NMO, demonstrated FMN-dependent oxidation of propionate 3-nitronate (P3N), and defined four characteristic sequence motifs^5^.

The subsequent characterization of PA1024 provides a direct warning against equating sequence annotation with enzymatic identity. Although originally annotated as a 2-nitropropane dioxygenase/NMO, purified PA1024 showed no detectable activity towards P3N, 3-NPA or several related nitro compounds and instead functioned predominantly as an NADH:quinone reductase^6^. Hundreds of PA1024-like proteins carrying related motifs had likewise been annotated as NMOs^6^. This precedent is highly relevant to EWG9 OXR01. Its strong COG2070/YrpB correspondence, substantial PF03060/NMO and NPD-like support, NMO-associated sequence motifs and phylogenetic placement establish membership within an overlapping YrpB/NMO-associated sequence space, but none independently demonstrates NMO catalysis. The observation that experimentally validated PA4202 itself produces a YrpB-dominant conserved-domain profile in our analysis further emphasizes that dominant domain correspondence cannot serve as a binary discriminator between NMO and non-NMO functions.

The biochemical diversity of nitrocompound-active oxidoreductases provides a second context for interpreting this sequence space. *Streptomyces ansochromogenes* NaoA shares similarity with proteins historically assigned as 2-nitropropane dioxygenases, including a *Bacillus subtilis* YrpB-related protein, yet the purified enzyme preferentially oxidizes nitroethane and 1-nitropropane and shows much weaker activity towards 2-nitropropane^42^. Flavin-dependent nitrocompound transformations can also proceed through fundamentally different redox chemistries; NAD(P)H:FMN oxidoreductase-mediated TNT reduction, for example, proceeds through reductive intermediates and can lead to enzyme inactivation^43^. These examples indicate that related flavoprotein architectures can support different substrates and reaction chemistries. The YrpB-dominated environmental reservoir detected here may therefore represent biochemical diversity rather than a collection of cryptic canonical NMOs.

The *Eisenia fetida* feed–gut–cast continuum provides an ecological bridge between this environmental protein diversity and cultivable phenotypes. Microbial detoxification of plant secondary metabolites is increasingly recognized across soil, rumen and animal-associated microbiomes, where toxins can become substrates for transformation or nutrient acquisition^44^. Particularly relevant parallels have emerged from herbivorous insects. The microbiota of *Nezara viridula* contributes to detoxification of plant-derived 3-NPA, and a gut-associated *Serratia* strain was shown to deplete NPA with concomitant accumulation of nitrate and nitrite^45^. A subsequent analysis found variable NPA tolerance among community members and demonstrated detoxification in six of eight tested isolates, reinforcing the idea that a common toxin-responsive phenotype can be distributed across taxonomically and physiologically distinct bacteria^46^. Our recovery of 31 3-NPA-responsive isolates representing 24 genera from the earthworm-associated continuum extends this pattern to a soil– detritivore system.

EWG9 provided a tractable strain-level example within this diversity. It combined rapid and sustained growth with measurable parent 3-NPA depletion and remained viable across a tenfold concentration range. Importantly, its strongest proliferation occurred when glucose supplied carbon and 3-NPA replaced nitrate as the nitrogen-containing substrate, whereas growth was weaker when 3-NPA was supplied as the sole carbon source. This phenotype suggests that EWG9 handles 3-NPA most effectively when energetic carbon requirements are met independently rather than behaving as a specialist that efficiently uses 3-NPA as a stand-alone carbon substrate. The approximately 44% disappearance of parent 3-NPA over 36 h, together with the stability of abiotic and *E. coli* controls, establishes a biological depletion phenotype. It does not, however, identify the transformation products or demonstrate mineralization. This is an important distinction from systems in which nitrite, nitrate, carbonyl products or other metabolites have been directly resolved ^42,45^.

The composite transcriptome suggests that the EWG9 phenotype is accompanied by broader physiological adjustment rather than strong activation of a single candidate gene. Increased replication/translation-associated expression occurred together with shifts in glycine/C1 metabolism, respiratory/redox functions and carbon allocation, whereas canonical nitrate/nitrite-associated functions were not coordinately induced. Most notably, the OXR01 locus itself changed only marginally. A similarly broad metabolic dimension has been observed in *N. viridula*-associated bacteria, where NPA exposure was associated with perturbation of amino-acid metabolism and transport, including possible effects on leucine biosynthesis^46^. These observations leave several possibilities open for EWG9, including constitutive OXR01 expression, transient induction outside the sampled window, post-transcriptional control, or participation of additional enzymes. Because the RNA-seq samples were pooled before extraction, the transcriptome remains a condition-level response map rather than causal evidence for OXR01 function.

Sequence conservation and structural modelling narrow this uncertainty but do not resolve it. Class I NMO active-site studies have shown that even conserved residues are not simple determinants of activity: replacement of several active-site tyrosines and Lys307 in PA4202 caused relatively modest changes in kinetic parameters, suggesting distributed contributions to substrate recognition rather than dependence on a single catalytic signature^25^. Conversely, recent characterization of the *Psychrobacter* PsNMO illustrates the evidentiary threshold required for functional assignment: the gene was cloned, the recombinant enzyme purified, enzymatic activity measured and product formation demonstrated^22^. Against this standard, docking, hydrogen-bond analysis, 300-ns molecular dynamics and pocket characterization show that EWG9 OXR01 is structurally compatible with 3-NPA, but cannot establish catalytic turnover.

The broader implication is therefore not that YrpB should replace NMO as an annotation for 3-NPA metabolism, but that the environmental YrpB/NMO-associated repertoire defines a functional search space that remains substantially underexplored. Microbial degradation of plant toxins frequently evolves through different enzymes and even distinct pathways converging on the same ecological phenotype^44^. The YrpB-dominated sequence reservoir identified here may similarly contain proteins with canonical NMO activity, alternative nitrocompound redox activities, indirect roles in cellular detoxification, or functions unrelated to nitrotoxins. Distinguishing these possibilities will require purified-protein kinetics, reaction-product identification and genetic loss- and gain-of-function experiments. Within that broader repertoire, EWG9 OXR01 currently remains an NMO-associated YrpB-family oxidoreductase with unresolved catalytic identity. Future biochemical evidence may show that it performs another YrpB-family redox function—or it may ultimately establish EWG9 OXR01 as a previously unrecognized, divergent bacterial NMO. That possibility is precisely what makes this broader microbial nitrotoxin-associated oxidoreductase space worth examining.

## SUPPLEMENTARY FIGURES AND TABLES

### Supplementary Figures

**Supplementary Figure 1:**
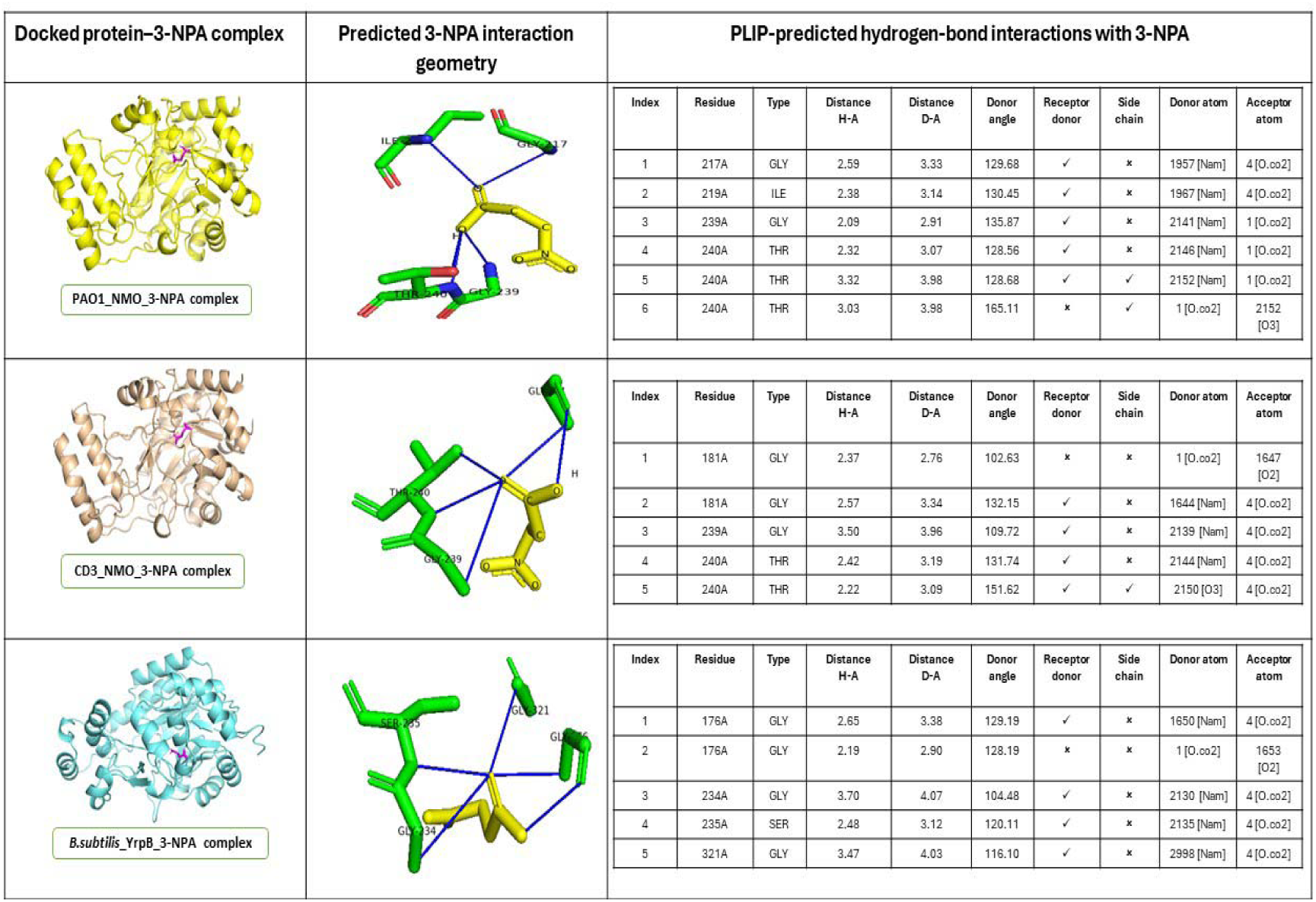
Comparative PLIP analysis of hydrogen-bond interactions in 3-NPA-bound NMO/YrpB complexes. Docked 3-NPA complexes of *Pseudomonas aeruginosa* PAO1 NMO, *P. aeruginosa* CD3 NMO and *Bacillus subtilis* YrpB are shown in the left panels. Middle panels depict the predicted 3-NPA interaction geometry, with hydrogen-bond contacts indicated by blue lines. Right panels summarize PLIP-predicted hydrogen-bond interactions, including interacting residues, hydrogen–acceptor (H–A) and donor–acceptor (D–A) distances, donor angles, receptor-donor status, side-chain involvement and donor/acceptor atoms. PAO1 NMO formed six predicted hydrogen bonds with 3-NPA, whereas CD3 NMO and *B. subtilis* YrpB each formed five. These interactions describe predicted contact architecture and should not be interpreted as direct measures of binding affinity or catalytic activity.

**Supplementary Figure 2:**
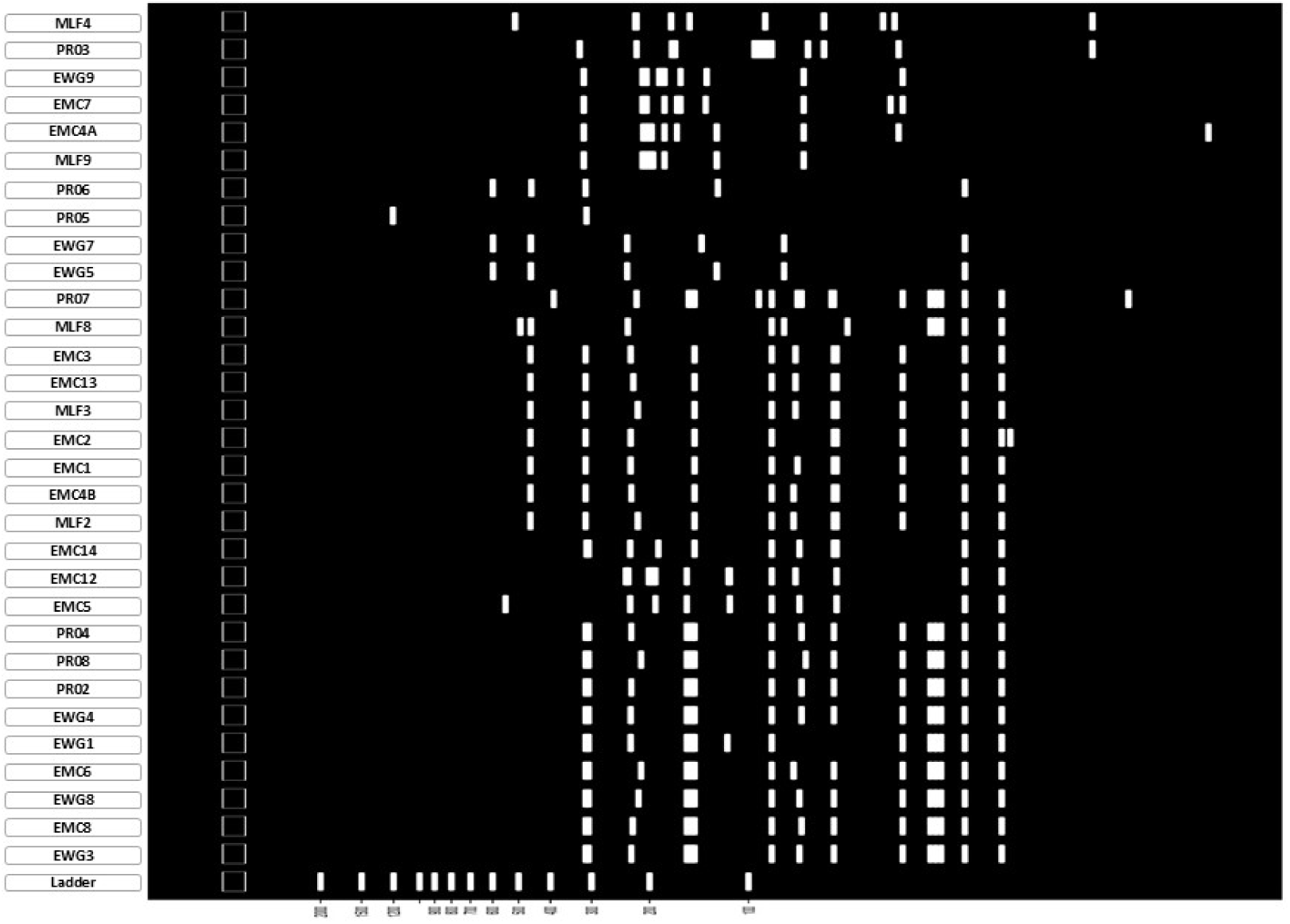
Virtual HaeIII restriction profiles of 16S rRNA gene amplicons from cultivable 3-NPA-responsive isolates. In silico HaeIII digestion profiles are shown for the 31 isolates recovered from the *Eisenia fetida* feed– gut–cast continuum. White bands indicate predicted restriction fragments, with the 100-bp DNA ladder shown at the bottom for size reference. Isolate prefixes denote source: MLF, formulated feed; EWG, whole gut of formulated-feed-fed earthworms; EMC, casts from formulated-feed-fed earthworms; and PR, isolates from the corresponding normal-feed control continuum. The virtual restriction patterns provide a complementary fingerprint of sequence-level diversity among the cultivable isolates and are interpreted alongside, rather than independently of, the 16S rRNA gene phylogeny.

**Supplementary Figure 3:**
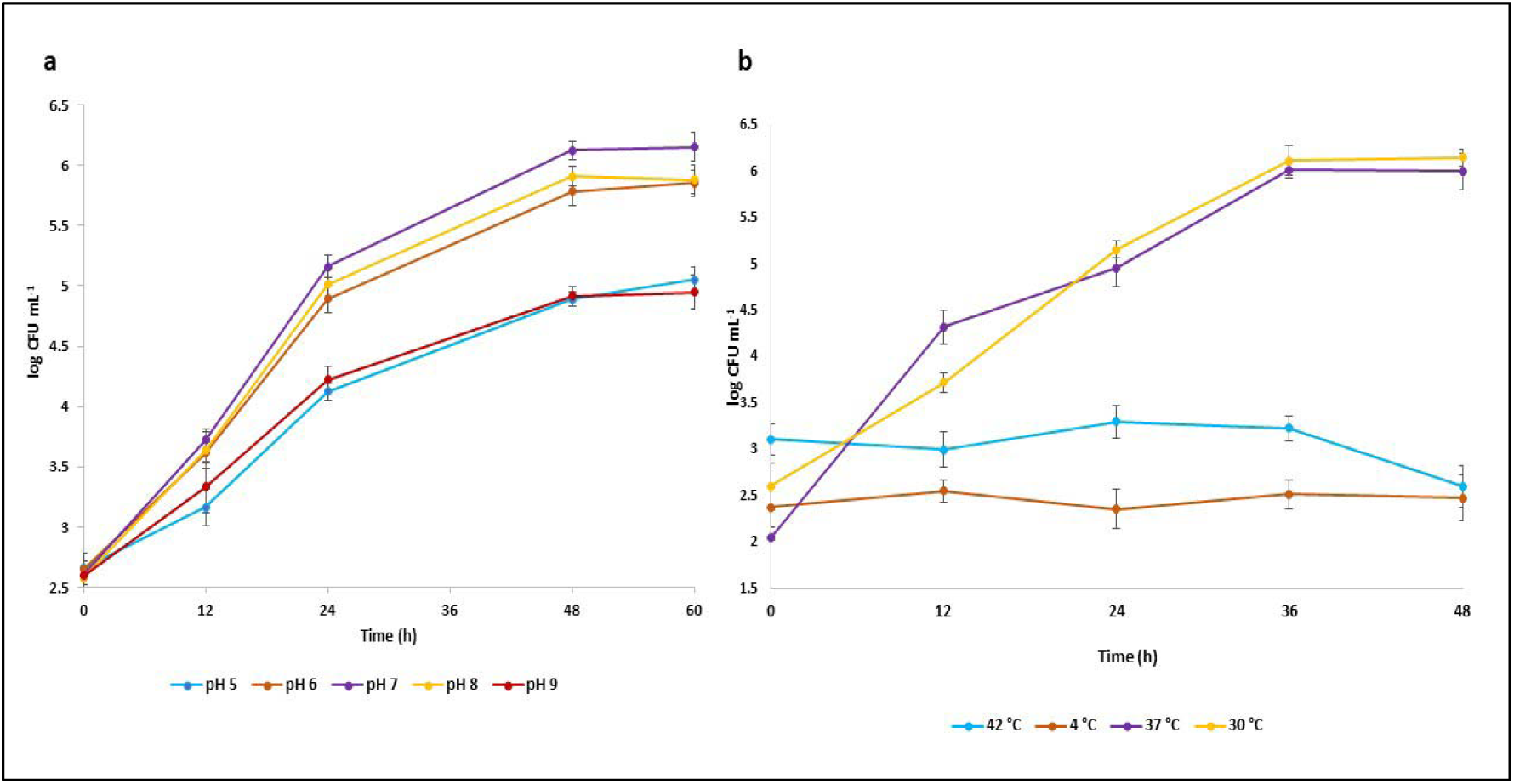
Effects of pH and temperature on growth of *Serratia* sp. EWG9 in 3-NPA-containing medium. **a,** Growth of EWG9 at pH 5, 6, 7, 8 and 9 over 60 h, expressed as log CFU ml□¹. The strongest growth was observed near neutral to mildly alkaline conditions, particularly at pH 7–8, whereas growth was lower at pH 5, 6 and 9. **b,** Growth of EWG9 at 4, 30, 37 and 42 °C over 48 h. Robust growth occurred at 30 and 37 °C, while little or no sustained proliferation was observed at 4 or 42 °C. Data are presented as mean values with error bars representing variability among replicates.

**Supplementary Figure 4:**
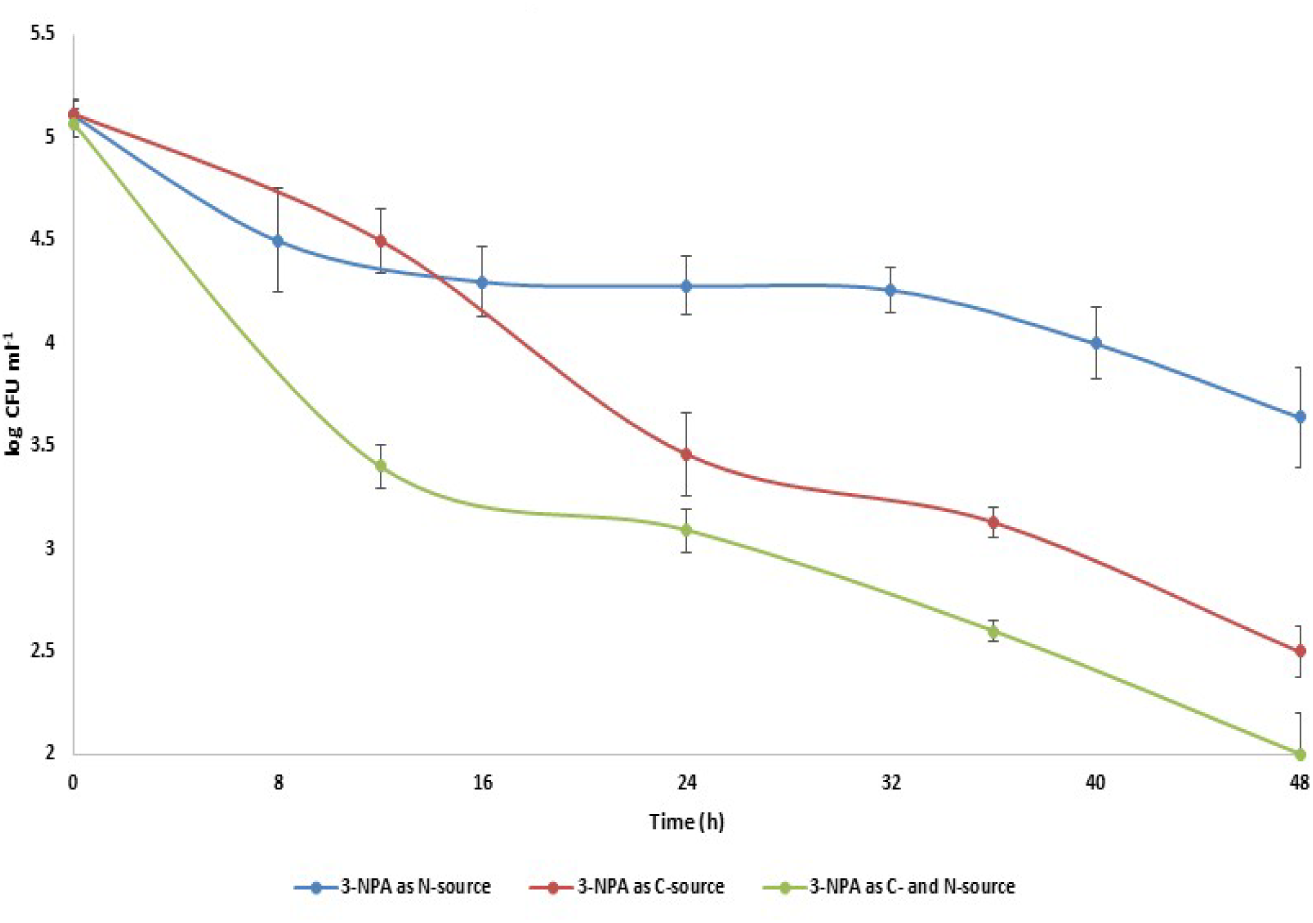
Growth of *Escherichia coli* K-12 under alternative 3-NPA nutritional conditions. Viable-cell counts of *E. coli* K-12 were monitored for 48 h in mineral salts medium containing 3-NPA as the sole nitrogen source, sole carbon source or combined carbon and nitrogen source. Growth declined under all three conditions, with the greatest loss of viability when 3-NPA served as the combined carbon and nitrogen source and the least decline when it served only as the nitrogen source. Points represent mean logource. Points represent m□ CFU ml□¹ and error bars indicate SD (*n* = 3 biological replicates). Two-way mixed-design repeated-measures ANOVA using the common 0-, 24- and 48-h time points showed a significant effect of time (*F□*,□□= 419.40, *P* < 0.001, ηp² = 0.986) and a significant time × condition interaction (*F□*,□□= 24.67, *P* < 0.001, ηp² = 0.892), indicating that the decline in viable counts differed among the three nutritional configurations.

**Supplementary Figure 5:**
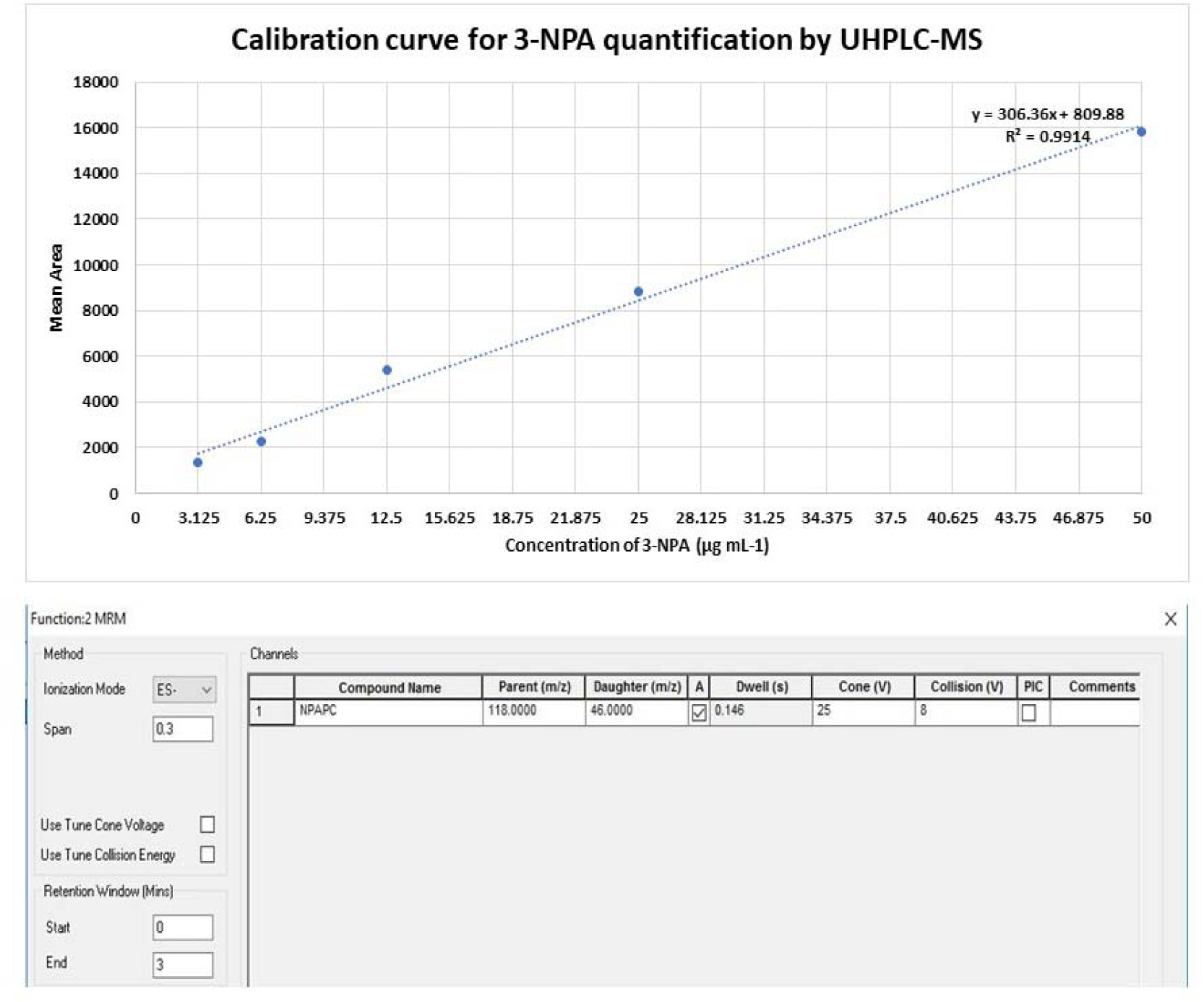
UHPLC–MS/MS calibration and MRM settings used for 3-NPA quantification. **a,** Calibration curve generated from 3-NPA standards spanning 3.125–50 µg ml□¹, showing a linear response between concentration and mean peak area (y = 306.36x + 809.88; R² = 0.9914). **b,** Multiple-reaction monitoring (MRM) parameters used for 3-NPA detection in negative electrospray ionization mode (ESI−), with the monitored transition m/z 118.0 → 46.0, dwell time 0.146 s, cone voltage 25 V and collision energy 8 V.

**Supplementary Figure 6:**
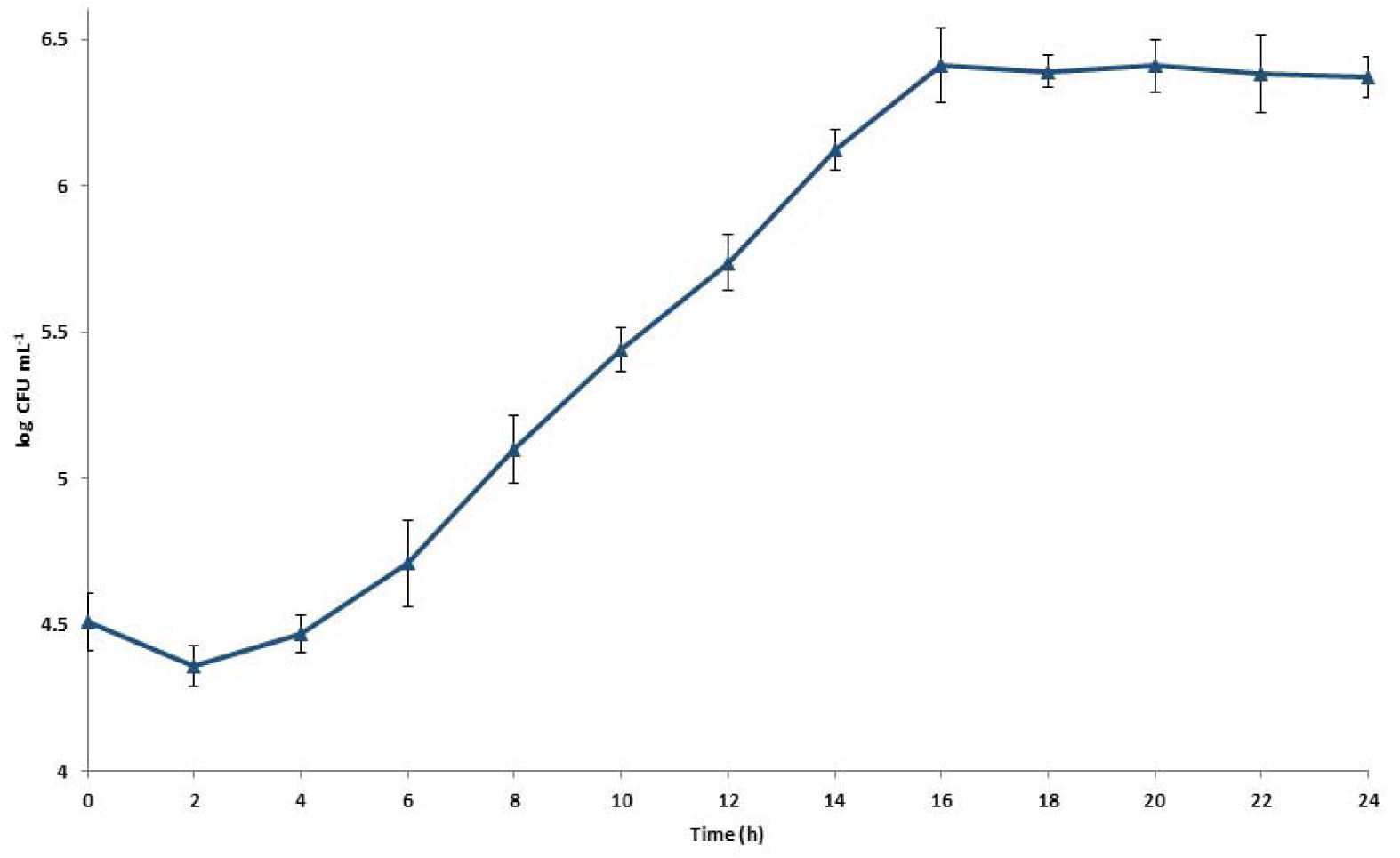
Growth kinetics of *Serratia* sp. EWG9 with 3-NPA as the sole nitrogen source. Viable-cell density of *Serratia* sp. EWG9 was monitored at 2-h intervals over 24 h in glucose-containing mineral salts medium in which 3-NPA served as the sole nitrogen source. Cell density increased from approximately 4.5 log□□ CFU ml□¹ at inoculation to approximately 6.4 log□□ CFU ml□¹ by 16 h, followed by a near-stationary phase through 24 h. Data are mean ± SD (*n* = 3 biological replicates). An AR(1) repeated-measures model showed a significant temporal increase in viable-cell density (*F□*,□.□□= 117.16, *P* < 0.001), whereas the quadratic time effect was not significant (*P* = 0.105). Within-culture observations showed significant first-order autocorrelation (ρ = 0.607, *P* < 0.001).

### Supplementary Table Legends

Supplementary Table 1: Comparative AutoDock Vina docking scores of 3-NPA and related nitro-containing ligands against NMO/YrpB-family oxidoreductases

Supplementary Table 2: Metadata and accession information for the 13 metagenomic datasets included in the comparative analysis.

Supplementary Table 3: Taxonomic identity, isolation source and GenBank accession numbers of the 31 cultivable 3-NPA-responsive bacterial isolates recovered from the earthworm feed–gut–cast continuum.

Supplementary Table 4: Correspondence between cultivable 3-NPA-responsive bacterial genera and their detection across amplicon-metagenomic datasets.

Supplementary Table 5: Inventory of *Eisenia fetida* gut-derived isolates, MAGs, and metagenomic assemblies carrying YrpB- or nitronate monooxygenase-family oxidoreductases.

## Declarations

### Ethical Approval

Ethical approval (IAEC/NBU/2022-34) for experimentation on earthworms was taken from the Institutional Animal Ethics Committee, under the guidelines of CPCSEA, New Delhi, India.

## Acknowledgements

The authors thank the Department of Biotechnology, University of North Bengal, and the Centre for Floriculture and Agri-Business Management (COFAM) for providing infrastructural support during the work. Authors are also grateful to Mr. Arup Ghosh for his assistance throughout the work. The authors also acknowledge the use of ChatGPT (GPT-5.0; OpenAI) during manuscript preparation, to assist in drafting narrative text. No AI tool was used to generate, manipulate, or analyze experimental data or results. All AI-generated content was critically reviewed to ensure accuracy and scientific integrity.

## Author Contributions

**Conceptualization:** Ranadhir Chakraborty.

**Data curation:** Partha Barman, Shilpa Sinha.

**Formal analysis:** Partha Barman, Shilpa Sinha, Ranadhir Chakraborty.

**Investigation:** Partha Barman, Shilpa Sinha.

**Methodology:** Partha Barman, Shilpa Sinha, Ranadhir Chakraborty.

**Resources:** Ranadhir Chakraborty.

**Validation:** Partha Barman.

**Writing – original draft:** Partha Barman, Shilpa Sinha, Ranadhir Chakraborty.

**Writing – review and editing:** All authors.

**Supervision and Correspondence:** Ranadhir Chakraborty.

## Data availability

The 16S rRNA gene sequences of the 31 cultivable 3-NPA-responsive bacterial isolates from this study have been deposited in GenBank under accession IDs PV648949, PV648974, PV648979, PV648998, PV649632, PV649633, PV649646, PV649860, PV649862, PV649869, PV650256, PV650259, PV650262, PV650263, PV650264, PV650265, PV650266, PV650289, MW205826, PV650303, PV650304, PV650351, PV650400, PV650402, PV650443, PV650611, PQ394639, PV650612, PV650613, PV650623 and PV650640. The RNA-sequencing data for *Serratia* sp. EWG9 are available through the NCBI Sequence Read Archive. Transcriptome sequencing of EWG9 grown in minimal salts medium with 3-nitropropionic acid as the sole nitrogen source is deposited under BioProject PRJNA821440, BioSample SAMN27097574 and SRA accession SRR18573213. The corresponding transcriptome generated with potassium nitrate as the sole nitrogen source is deposited under BioProject PRJNA821437, BioSample SAMN27094077 and SRA accession SRR18573214. The previously reported genome assembly of *Serratia* sp. EWG9 is available in DDBJ/ENA/GenBank under accession JADNRO000000000. Accession information for the 13 publicly available metagenomic datasets analysed in this study is provided in Supplementary Table 2.

