## Supplemental Table 1 for "Nitrotoxin metabolism in bacteria may have emerged from a diverse oxidoreductase reservoir"

| **Ligand** | ***Serratia sp.* EWG9 YrpB** | ***P. aeruginosa* PAO1 NMO** | ***P. aeruginosa* CD3 NMO** | ***B. subtilis* YrpB** |
| --- | --- | --- | --- | --- |
| **3-NPA** | -5.3 | -5.0 | -5.1 | -4.9 |
| Nitromethane | -3.3 | -3.3 | -3.3 | -3.2 |
| Nitroethane | -3.7 | -3.7 | -3.7 | -3.6 |
| 1-Nitropropane | -4.1 | -4.1 | -4.1 | -4.0 |
| 2-Nitropropane | -4.0 | -4.1 | -4.1 | -4.2 |
| Chloropicrin | -4.4 | -4.2 | -4.2 | -4.3 |
| Tetranitromethane | -5.6 | -5.7 | -5.9 | -6.1 |
| 3-Nitropropanol | -4.4 | -4.4 | -4.4 | -4.1 |
| Miserotoxin | -7.3 | -7.0 | -6.0 | -6.9 |
| 2-Nitro-2-butene | -4.6 | -4.6 | -4.5 | -4.7 |
| TNT | -6.9 | -6.9 | -6.9 | -7.1 |
| Tetryl | -5.7 | -5.2 | -5.5 | -5.5 |
| Nitroglycerin | -6.4 | -7.0 | -7.0 | -6.2 |
| RDX | -7.1 | -7.7 | -7.5 | -6.7 |

**Supplementary Table 1 : Comparative AutoDock Vina docking scores of 3-NPA and related nitro-containing ligands against NMO/YrpB-family oxidoreductases**

Docking scores are reported in kcal mol^−1^ for the top-ranked pose obtained for each protein–ligand pair

Notes. More negative AutoDock Vina scores indicate more favourable predicted docking according to the Vina scoring function; they are comparative docking scores and should not be interpreted as experimentally measured binding free energies.

Blind docking used a uniform 40 × 40 × 40 Å³ search space centred individually on each receptor, with the top-ranked pose retained for comparison.

Abbreviations: 3-NPA, 3-nitropropionic acid; TNT, 2,4,6-trinitrotoluene; tetryl, trinitrophenylmethylnitramine; RDX, cyclotrimethylenetrinitramine (hexogen); NMO, nitronate monooxygenase.
