## Supplemental Table 3 for "Nitrotoxin metabolism in bacteria may have emerged from a diverse oxidoreductase reservoir"

| **STRAIN** | **ORGANISM** | **Isolation source** | **GenBank Accession ID** |
| --- | --- | --- | --- |
| **PR02** | *Cellulosimicrobium composti* | Control feed (PrCD)-fed earthworm microbe system | PV648949 |
| **PR03** | *Acinetobacter indicus* | Control feed (PrCD)-fed earthworm microbe system | PV648974 |
| **PR04** | *Cellulosimicrobium cellulans* | Control feed (PrCD)-fed earthworm microbe system | PV648979 |
| **PR05** | *Staphylococcus equorum* subsp. *linens* | Control feed (PrCD)-fed earthworm microbe system | PV648998 |
| **PR06** | *Bacillus xiamenensis* | Control feed (PrCD)-fed earthworm microbe system | PV649632 |
| **PR07** | *Rhodococcus parequi* | Control feed (PrCD)-fed earthworm microbe system | PV649633 |
| **PR08** | *Cellulosimicrobium cellulans* | Control feed (PrCD)-fed earthworm microbe system | PV649646 |
| **MLF2** | *Glutamicibacter mysorens* | Formulated feed | PV649860 |
| **MLF3** | *Glutamicibacter nicotianae* | Formulated feed | PV649862 |
| **MLF4** | *Stenotrophomonas maltophilia* | Formulated feed | PV649869 |
| **MLF8** | *Brevibacterium sediminis* | Formulated feed | PV650256 |
| **MLF9** | *Leclercia barmai* | Formulated feed | PV650259 |
| **EWG1** | *Cellulosimicrobium sp.* | Formulated feed-fed earthworm gut | PV650262 |
| **EWG3** | *Cellulosimicrobium cellulans* | Formulated feed-fed earthworm gut | PV650263 |
| **EWG4** | *Cellulosimicrobium cellulans* | Formulated feed-fed earthworm gut | PV650264 |
| **EWG5** | *Priestia aryabhattai* | Formulated feed-fed earthworm gut | PV650265 |
| **EWG7** | *Priestia aryabhattai* | Formulated feed-fed earthworm gut | PV650266 |
| **EWG8** | *Cellulosimicrobium cellulans* | Formulated feed-fed earthworm gut | PV650289 |
| **EWG9** | *Serratia* sp. | Formulated feed-fed earthworm gut | MW205826 |
| **EMC1** | *Glutamicibacter nicotianae* | Castings from formulated feed-fed earthworms | PV650303 |
| **EMC2** | *Glutamicibacter nicotianae* | Castings from formulated feed-fed earthworms | PV650304 |
| **EMC3** | *Glutamicibacter nicotianae* | Castings from formulated feed-fed earthworms | PV650351 |
| **EMC4A** | *Leclercia barmai* | Castings from formulated feed-fed earthworms | PV650400 |
| **EMC4B** | *Glutamicibacter mysorens* | Castings from formulated feed-fed earthworms | PV650402 |
| **EMC5** | *Microbacterium ihumii* | Castings from formulated feed-fed earthworms | PV650443 |
| **EMC6** | *Cellulosimicrobium cellulans* | Castings from formulated feed-fed earthworms | PV650611 |
| **EMC7** | *Leclercia barmai* | Castings from formulated feed-fed earthworms | PQ394639 |
| **EMC8** | *Cellulosimicrobium composti* | Castings from formulated feed-fed earthworms | PV650612 |
| **EMC12** | *Microbacterium paraoxydans* | Castings from formulated feed-fed earthworms | PV650613 |
| **EMC13** | *Glutamicibacter nicotianae* | Castings from formulated feed-fed earthworms | PV650623 |
| **EMC14** | *Brachybacterium rhamnosum* | Castings from formulated feed-fed earthworms | PV650640 |

**Supplementary Table 3: Taxonomic identity, isolation source and GenBank accession numbers of the 31 cultivable 3-NPA-responsive bacterial isolates recovered from the earthworm feed–gut–cast continuum.**
